# Constitutive expression of CX3CR1-BAC-Cre introduces minimal off-target effects in microglia

**DOI:** 10.1101/2024.11.01.621625

**Authors:** Fadya H. Mroue-Ruiz, Bhoomi Desai, Madison Garvin, Jonila Shehu, Faith Kamau, Urjoshi Kar, Jessica L. Bolton

## Abstract

CX3CR1-Cre mouse lines have produced important advancements in our understanding of microglial biology. Recent studies have demonstrated the adverse effects of tamoxifen-induced CX3CR1-Cre expression during development, which include changes in microglial density, phenotype, and DNA damage, as well as anxiety-like behavior. However, the unintended effects of constitutive CX3CR1-BAC-Cre expression remain unexplored. Here, we characterized the effects of CX3CR1-BAC-Cre expression on microglia in CX3CR1-BAC-Cre+/- and CX3CR1-BAC-Cre-/-male and female littermates during early postnatal development and adulthood in multiple brain regions. Additionally, we performed anxiety-like behavior tests to assess changes caused by Cre expression. We found that CX3CR1-BAC-Cre expression causes subtle region- and sex-specific changes in microglial density, volume, and morphology during development, but these changes normalized by adulthood in all brain regions except the hippocampus. No behavioral effects were found. Our findings suggest that the constitutive-Cre model might be less detrimental than the inducible model, and highlight the need for proper controls.

## INTRODUCTION

The Cre-lox system is a widely used gene-editing tool discovered in the bacteriophage P1^1^. To function, the system needs the presence of both the Cre-enzyme and specific DNA sequences, called loxP. When together, the Cre enzyme is able to recognize and cleave the loxP sites, allowing for precise genetic modifications. Currently, it is possible to produce cell-specific transgenic animals by adding the Cre-driver under a cell-specific promoter. Furthermore, the Cre-lox system provides a choice of two models for genetic manipulation with high spatial specificity: inducible and constitutive. The inducible Cre-lox requires use of an exogenous inducer like tamoxifen to induce Cre-recombinase expression, thus allowing for excellent temporal specificity over gene manipulation. In contrast, the constitutive model allows for constant Cre-recombinase expression throughout an animal’s life.

This system has been particularly important in the field of microglial biology. Microglia are known as the resident immune cells of the brain, although they have various roles in homeostasis, such as synaptic pruning in development^2^. Within the CNS parenchyma, the expression of CX3CR1, a fractalkine receptor activated by its neuron-secreted ligand CX3CL1, is highly specific to microglia. Thus, many landmark studies have heavily relied on constitutive and inducible Cre-lox lines under the control of the microglia-specific promoter CX3CR1 to study and characterize the multi-faceted functions of microglia^3-6^. Although these investigations have significantly enhanced our understanding of microglia, more recent studies have demonstrated unintended adverse effects of using the Cre-lox system in microglia.

Specifically, the early postnatal induction of CX3CR1^CreER(Litt) 7^ was shown to have adverse effects on microglia^8^. The authors found a lower microglial density, an altered phenotype, upregulation of their phagocytic function, and DNA damage in the Cre+ microglia of the developing brain. While the irregular microglial number and phenotype had normalized by adulthood, these unintended and non-specific effects of CX3CR1-driven Cre-recombinase expression in early postnatal microglia caused the animals to display increased anxiety-like behavior in adulthood^8^. Furthermore, a separate study also reported a significant loss of homeostatic P2RY12 expression in microglia after the early postnatal induction of CX3CR1^CreER(Litt) 9^. Interestingly, these effects were shown to be limited only to Cre-recombinase expression under the CX3CR1 promoter and to its induction during early postnatal ages^8^.

However, the off-target effects of constitutive Cre expression with a bacterial artificial chromosome (BAC) recombination strategy under the promoter CX3CR1 remain unexplored. The widespread use of the CX3CR1 promoter in generating microglia-specific gene manipulation, along with the recently characterized detrimental effects of inducible Cre-recombinase expression under this promoter, warrant an urgent need to validate the constitutive CX3CR1-Cre transgenic lines. We hypothesized that due to its uninterrupted and sustained expression from embryonic stages through adulthood, constitutive CX3CR1-Cre expression would lead to altered microglial morphology and function. Thus, we performed an in-depth characterization of the effects of constitutive CX3CR1-BAC-Cre (from the GENSAT Transgenic Project) expression on early postnatal and adult microglia, as well as adult mouse behavior. Our results, along with recent studies, have important implications for the use of the CX3CR1-driven Cre-lox system to study microglia, especially during development.

## RESULTS

### Constitutive expression of Cx3cr1-BAC-Cre results in minimal changes in canonical microglial markers across the developmental brain

Tamoxifen-mediated Cre-recombinase expression under the promoter CX3CR1 (CX3CR1^CreER(Litt)^) has been shown to reduce IBA1+ microglia density in pups (P7-P9), while leading to increased IBA1+ microglia volume and Sholl intersections^8^. The dyshomeostatic effects of inducible Cre-recombinase expression under CX3CR1 (CX3CR1^YFP-CreER(Litt)^) has also been characterized by a significant loss of P2RY12 expression in pups (P15) microglia^9^. Thus, the effects of constitutive CX3CR1-Cre expression on microglia were assessed throughout the brain, using canonical microglial markers such as IBA1, P2RY12, and the phagocytic marker CD68 in pups (P7-P9).

#### Parietal cortex

The expression of constitutive CX3CR1-Cre in females caused a decrease in the number of IBA1+ microglia (interaction of Genotype x Sex, F_(1,32)_=4.242; p<0.05; Šídák p<0.05; Fig.1B), as well as a decrease in IBA1 volume (interaction of Genotype x Sex, F_(1,32)_=5.567; p<0.05; Šídák p<0.05; Fig.1C). However, the Sholl-based morphology analysis revealed no difference in segment volume, soma volume, or average process thickness. A decrease in volume was also found with the P2RY12 marker for Cre-positive animals (main effect of Genotype, F_(1,32)_=7.175; p<0.05; Fig.1K). Overall, this might indicate that females are more susceptible to the unintended effects of Cre in microglia; however, the pattern for the number of cells differs depending on which marker microglial marker is used.

**Figure 1.**
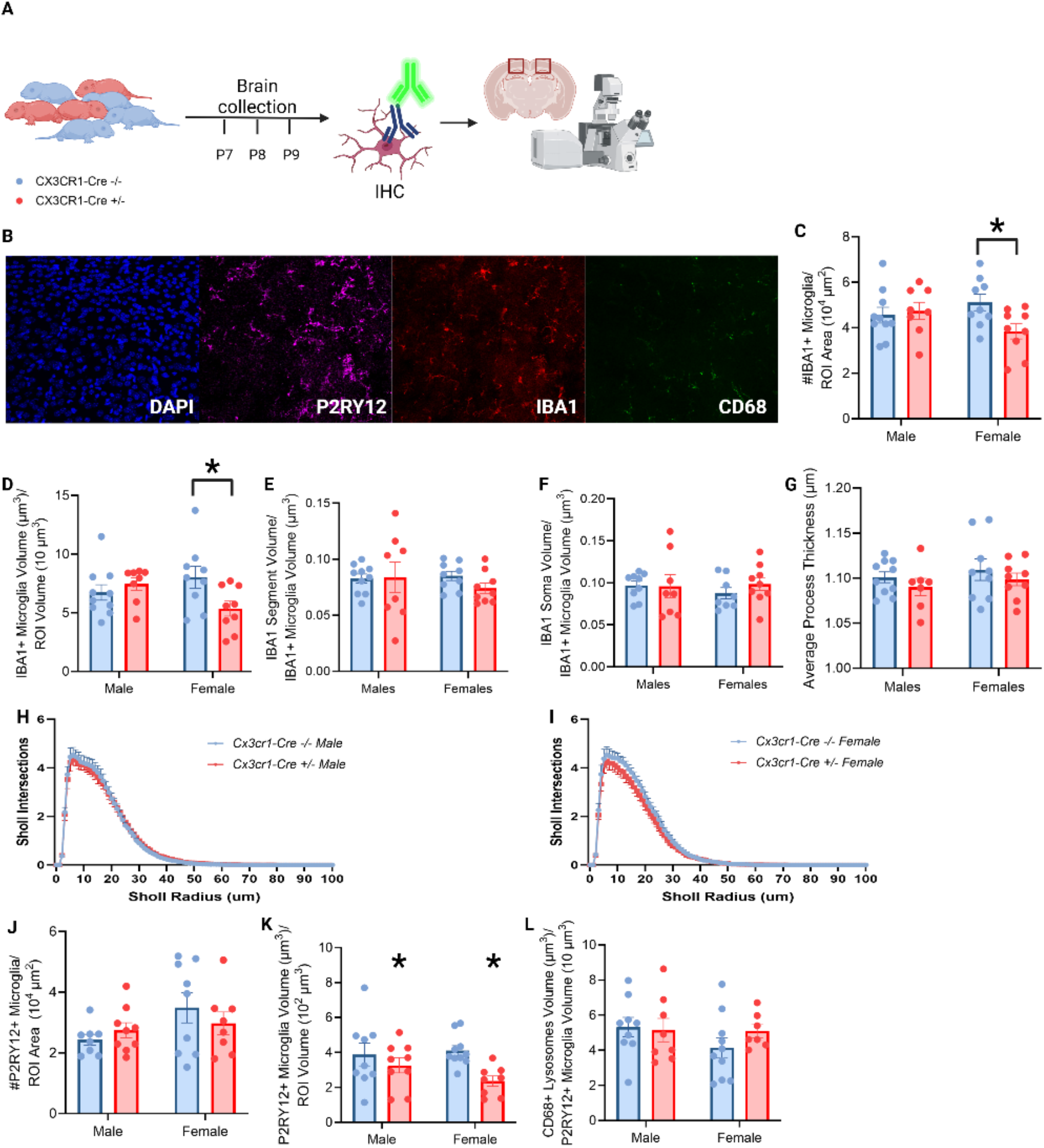
Constitutive CX3CR1-BAC-CRE expression does not induce major changes in the microglial phenotype during early postnatal development. A. Schematic of the experimental timeline. B. Representative images showing DAPI, P2RY12, IBA1 and CD68 staining in the PC. C. Quantification of IBA1+ microglia density (interaction of genotype x sex, F_(1,32)_=4.242; p<0.05; Šídák p<0.05). D. Quantification of IBA1+ microglia volume (interaction of genotype x sex, F_(1,32)=_5.567; p<0.05; Šídák p<0.05). E. (D-H) Sholl analysis metrics. Average segment volume of IBA1+ microglia. F. Average soma volume of IBA1+ microglia. G. Average process thickness of IBA1+ microglia. H. Number of Sholl intersections for each radius of IBA1+ microglia of male mice. I. Number of Sholl intersections for each radius of IBA1+ microglia of female mice. J. Quantification of P2RY12+ microglia density. K. Quantification of P2RY12+ microglia volume (main effect of genotype, F_(1,32)_=7.175; p<0.05). L. Quantification of CD68+ microglial lysosome volume relative to the P2RY12+ microglial volume. Each dot represents the average value for one animal. n=7-11. *p<0.05. Mean ± SEM. Schematic created using BioRender.

#### Paraventricular nucleus of the hypothalamus

No significant changes were found in the number or volume of IBA1+ microglia, but there was a trend for decreased volume in Cre+ animals (main effect of Genotype, F_(1,27)_=2.95, p<0.05, Fig.S1C), although there were no morphological differences based on this marker. The expression of Cre caused a significant decrease in the volume of P2RY12 (main effect of Genotype, F_(1,16)_=6.886, p<0.05; Fig.S1I) as well as increased complexity in the morphology of microglia in females, but not males (Interaction of Distance from soma x Genotype x Sex, F_(88,2904)_=1.812, p<0.05; 2-way ANOVA, F_(1,16)_=6.886, p<0.05, Fig.S1N). These results are in line with the PC, since females appear to be more susceptible to the expression of Cre in this particular line.

#### Amygdala

No changes were found in the P2RY12 density or volume. There was a significant increase in the total branch points of microglia in CX3CR1-Cre+ animals (main effect of Genotype, F_(1,32)_=7.582, p<0.05, Fig.S2D), as well as increased Sholl intersections in CX3CR1-Cre+ males (interaction of Distance from soma x Genotype x Sex, F_(102,3399)_=1.532, p<0.05; 2-way ANOVA, F_(1,16)_=9.000, p<0.05, Fig.S2F), suggesting that there are region- and sex-dependent differences. The CD68 lysosome volume remained unchanged.

#### CA1

The number of IBA1+ microglia was not affected by the expression of Cre. However, CX3CR1-Cre+ females showed decreased IBA1 volume (interaction of Genotype x Sex, F_(1,31)_=4.50, p<0.05, Šídák, p<0.05; Fig.S3C). No changes in any morphological metrics were found with either marker, but there was a trend for a decrease in P2RY12 microglial volume in Cre+ animals (main effect of Genotype, F_(1,35)_=3.33, p=0.07; Fig.S3I). Furthermore, the volume of CD68 lysosomes normalized to microglial volume was increased as a result of the expression of Cre (main effect of Genotype, F_(1,34)_=7.986, p<0.05; Fig.S3J), but this appears to be driven by the decrease in microglial volume (i.e., no significant difference was found in the raw CD68 volume).

#### CA3

The IBA1 volume was increased in Cre+ animals (main effect of Genotype, F_(1,31)_=6.28, p<0.05; Fig.S4C), but there were no changes in microglial morphology found in this brain region with either microglial marker. Additionally, females had a lower volume of CD68 lysosomes overall than males (main effect of Sex, F_(1,30)_=5.752, p<0.05; Fig.S4J).

#### Dentate Gyrus

No changes were found in the IBA1 number, volume, or metrics associated with microglial morphology. There was a trend for a decrease in the density of P2RY12+ microglia in Cre+ animals (main effect of Genotype, F_(1,32)_=3.26, p=0.0804, Fig.S5H).

### Microglial DNA is not damaged by the constitutive expression of CX3CR1-BAC-Cre

Microglia develop from their myeloid progenitors in the yolk sac at embryonic day (E)8.5, and they start to proliferate and colonize the CNS parenchyma before the blood-brain barrier is formed^10^. It has been shown that tamoxifen-mediated Cre-recombinase expression (CX3CR1^CreER(Litt)^) decreases the number of proliferating microglia, thereby suggesting decreased proliferation of early postnatal microglia, as well as induces DNA damage in the early postnatal period^8^. Thus, the effects of constitutive CX3CR1-BAC-Cre expression on proliferation and DNA damage were studied by immunostaining brain tissue with Ki67 (proliferating cells marker), IBA1 (microglial marker) and phosphorylated-γ-H2AX (DNA damage marker).

Interestingly, none of the tested brain regions showed changes due to genotype (Fig. 2 and Fig.S6-S9), indicating that constitutive CX3CR1-BAC-CRE expression is less detrimental to the cell cycle and DNA integrity in microglia compared to tamoxifen induced-Cre.

**Figure 2.**
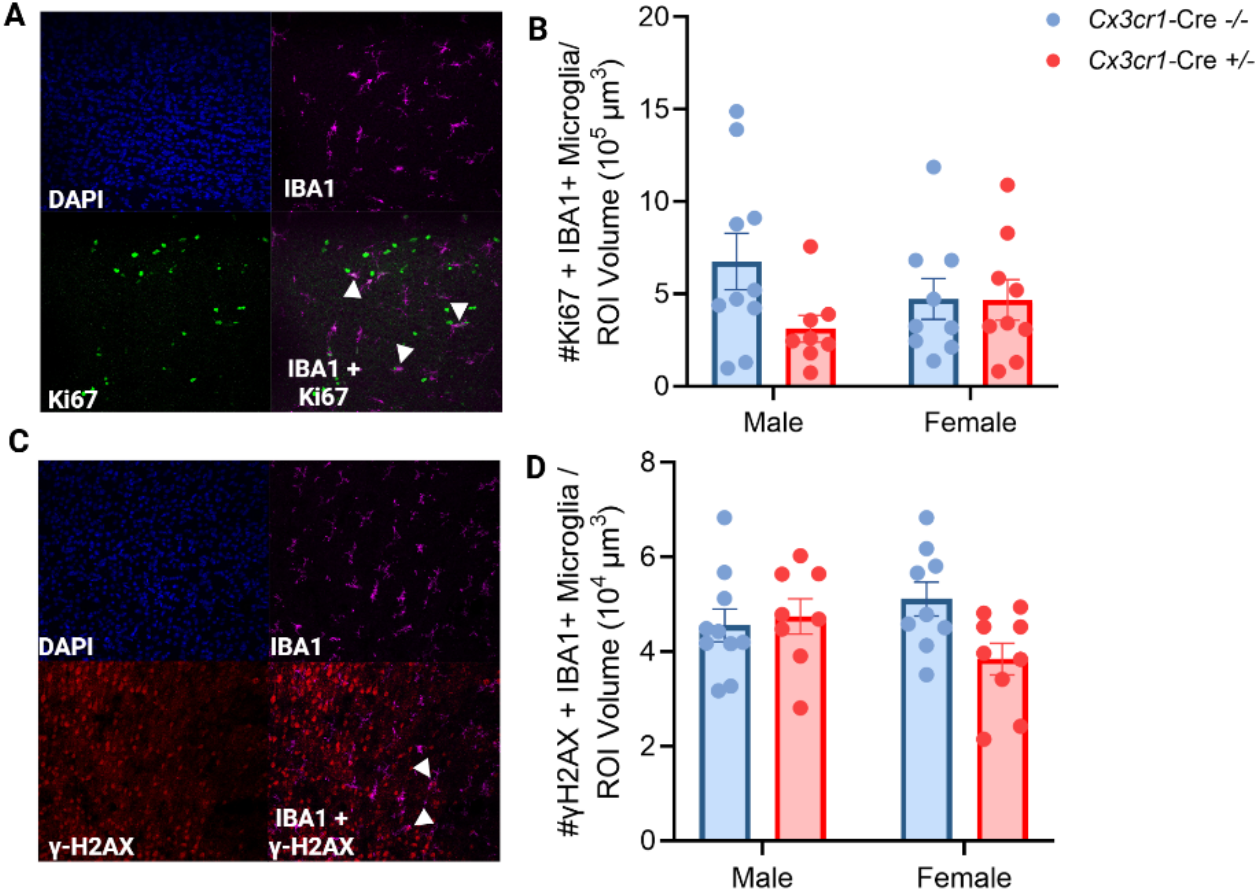
Microglial proliferation and DNA integrity during early postnatal development are not altered by CX3CR1-BAC-CRE expression. A. Representative images showing PC stained with DAPI, Ki67 and IBA1. White arrowheads point to a cell double-positive for Ki67 and IBA1. B. Quantification of density of cells double-positive for Ki67 and IBA1. C. Representative image showing a PVN stained with DAPI, IBA1 and phosphorylated-γ-H2AX. White arrowheads point to a cell double-positive for Ki67 and phosphorylated-γ-H2AX. D. Quantification of density of cells double-positive for phosphorylated-γ-H2AX and IBA1. Each dot represents the average value for one animal. n=7-9. Mean ± SEM.

**Figure 3.**
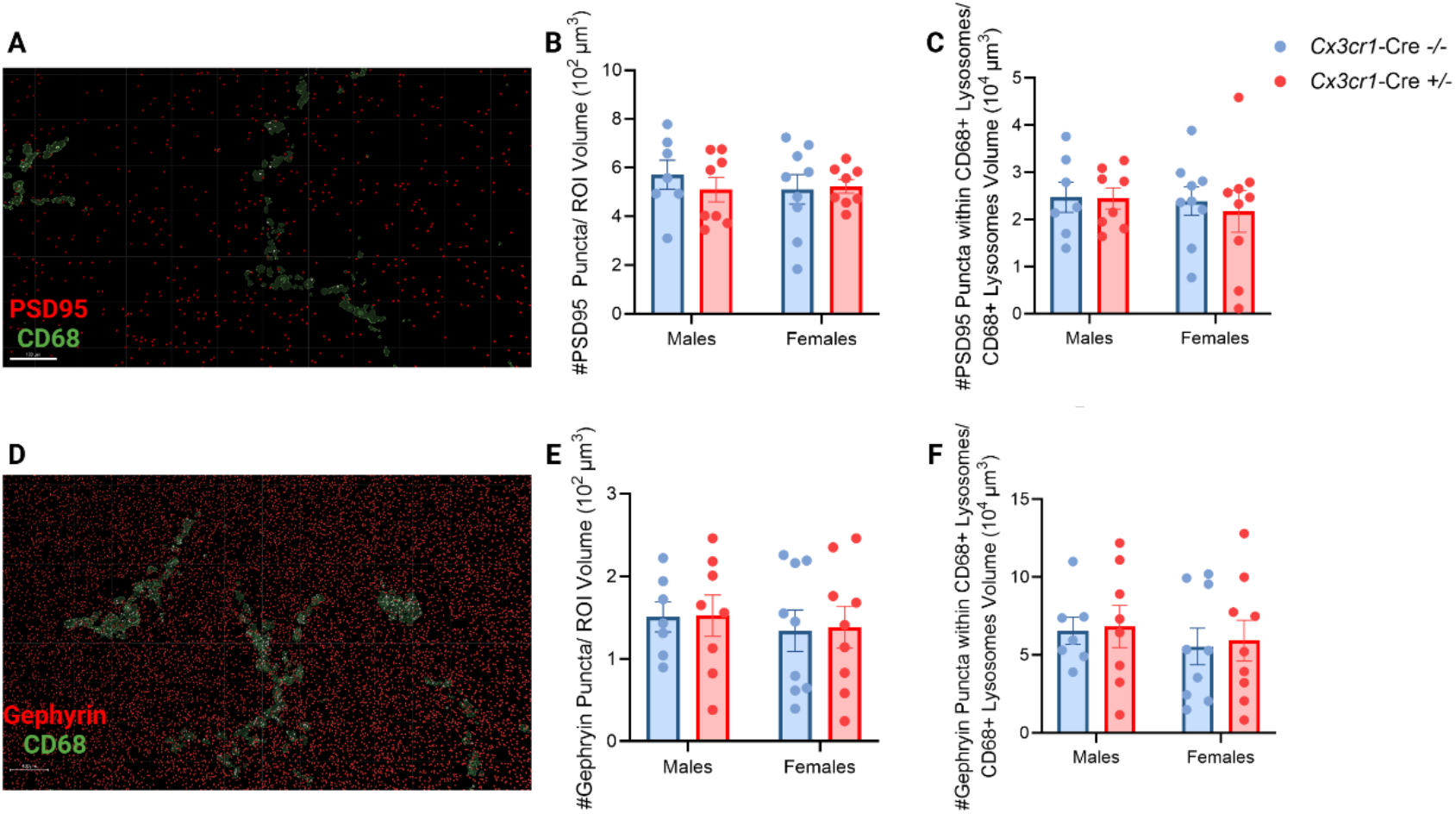
CX3CR1-BAC-CRE expression does not affect synaptic engulfment in the PC during early postnatal development. A. Representative image showing PSD95 puncta, and the reconstruction of CD68 volume in green. Non-colocalized PSD95 puncta in red, and colocalized puncta in white. B. Density quantification of PSD95 puncta. C. Density quantification of PSD95 puncta within CD68+ microglial lysosomes. D. Representative image showing gephyrin puncta, and the reconstruction of CD68 volume in green. Non-colocalized gephyrin puncta in red, and colocalized puncta in white. E. Density quantification of gephyrin puncta. F. Density quantification of gephyrin puncta within CD68+ microglial lysosomes. Each dot represents the average value for one animal. n=7-9. Mean ± SEM.

### Microglial engulfment of PSD95 and gephyrin is unchanged by CX3CR1-BAC-Cre expression in the developing brain, except in the CA1 of the hippocampus

Microglia are known to play a vital role during typical development in shaping neuronal circuitry by pruning synapses^2,11,12^. Furthermore, early postnatal tamoxifen-mediated induction of CX3CR1^CreER(Litt)^ has been shown to enhance microglial synaptic engulfment in pups^8^. Thus, microglial engulfment of post-synaptic structures during early postnatal development was assessed through immunostaining for PSD95 (excitatory post-synaptic terminal marker), gephyrin (inhibitory post-synaptic terminal marker), and CD68 (microglial lysosome marker) to test the effects of constitutive CX3CR1-BAC-Cre expression on microglial phagocytic function. Among the brain regions tested in this section of the study (PC, PVN, and CA1), only the CA1 showed an effect, with Cre expression resulting in decreased PSD95 puncta engulfment (main effect of Genotype, F_(1,29)_=4.453, p<0.05; Fig. S11C). In the same region, engulfment of gephyrin puncta within CD68 lysosomes was decreased as a result of constitutive Cre expression (main effect of Genotype F_(1,29)_=5.615, p<0.05; Fig. S11F), and females had less colocalization overall (main effect of Sex F_(1,29)_=6.670, p<0.05; Fig. S11F). Finally, there was a trend for a decrease in the density of gephyrin puncta in Cre+ animals (main effect of Genotype, F_(1,29)_=3.691, p=0.0646; Fig. S11E), which may suggest that the decrease in microglial engulfment was a compensatory mechanism. These region-specific results might suggest susceptibility of certain populations of microglia to the expression of Cre based on their spatial location in the brain.

#### The expression of CX3CR1-BAC-CRE does not affect the threat response or cause anxiety-like behavior in adulthood

It is widely known that alterations in microglia can lead to alterations in behavior in mice^5,13-15^. Furthermore, early postnatal tamoxifen-mediated induction of CX3CR1^CreER(Litt)^ has been shown to increase anxiety-like behavior in adulthood, with CX3CR1^CreER(Litt)^ animals that received tamoxifen spending significantly less time in the center of the open-field test (OFT) and in the open arms of the elevated-plus maze test (EPM)^8^. Thus, a battery of behavioral tests was performed in adult CX3CR1-BAC-Cre+/- and CX3CR1-BAC-Cre-/-mice (Fig. 4A).

**Figure 4.**
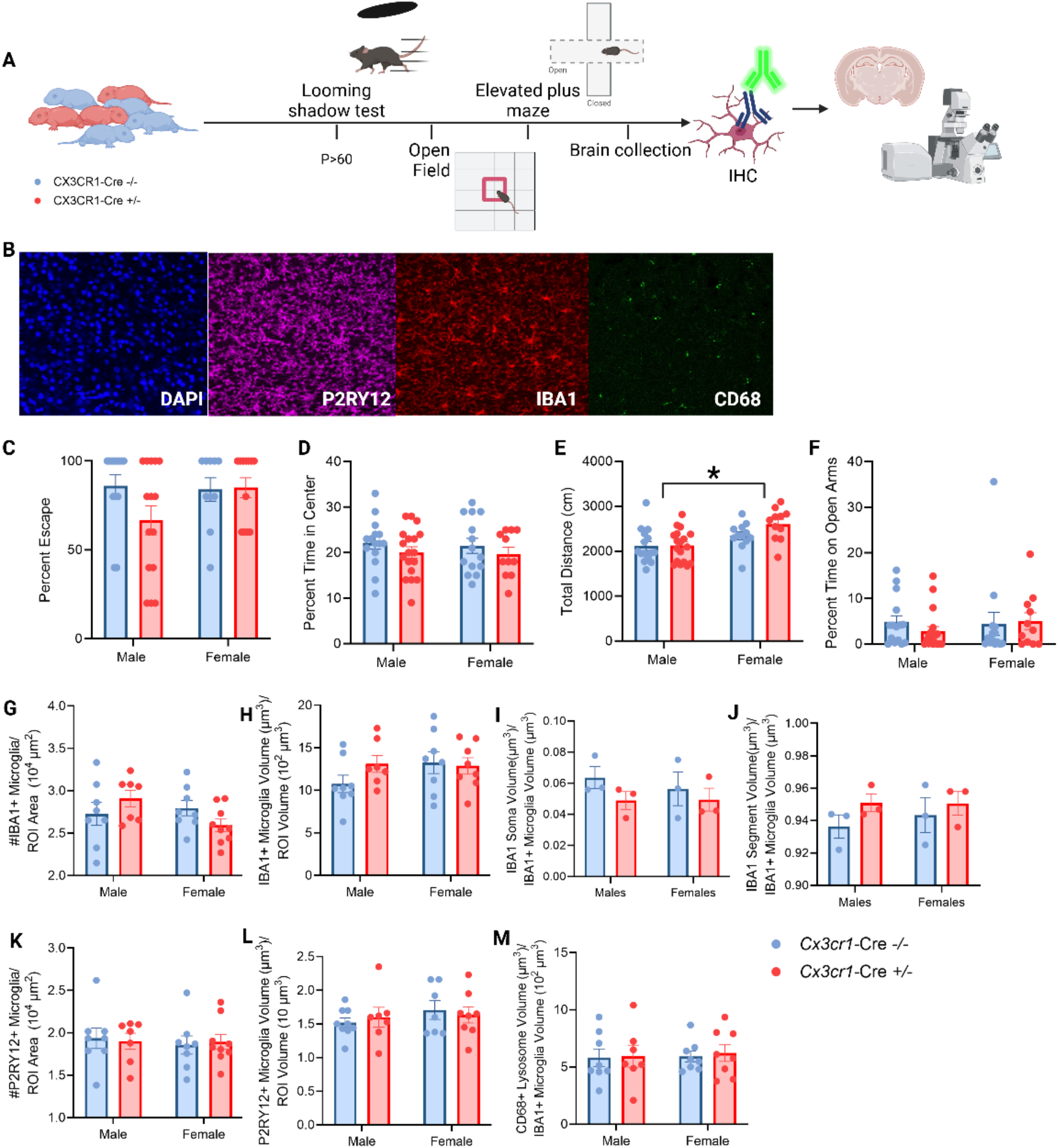
CX3CR1-BAC-CRE expression does not affect microglial phenotype or behavior in adulthood. A. Diagram representing the experimental timeline and order of the behavioral tasks. B. Representative images showing DAPI, P2RY12, IBA1 and CD68 staining in the PC. C. Quantification for the percent escape in the looming-shadow threat test. D. Quantification of the percent time spent by a mouse in the center of the open field. E. Quantification of total distance traveled by a mouse in the open field (main effect of genotype, F_(1,52)_=13.70, p<0.05). F. Quantification of the percentage time spent in the open arms of the elevated plus maze. n=10-15 animals/group. Each dot represents the average value for one animal. *p<0.05. G. Quantification of IBA1+ microglia density. H. Quantification of IBA1+ microglia volume. I. Average soma volume of IBA1+ microglia. J. Average segment volume of IBA1+ microglia. K. Quantification of P2RY12+ microglia density. L. Quantification of P2RY12+ microglia volume. M. Quantification of CD68+ microglial lysosome volume relative to the P2RY12+ microglial volume. Each dot represents the average value for one animal. n=3-8. *p<0.05. Mean ± SEM. Mean ± SEM. Schematic created using BioRender.

Unlike the inducible CX3CR1-Cre model, male and female adult mice with constitutive CX3CR1-BAC-Cre expression did not exhibit anxiety-like behavior in the OFT (Fig. 5D) or EPM (Fig. 5F), as there was no difference in the amount of time spent in the center of the open field or the amount of time spent in the open arms, respectively. The total distance travelled in the OFT was significantly higher in females as compared to males (main effect of Sex, F_(1,52)_=13.70, p<0.05; Fig.5E), but there were no significant differences due to genotype. Furthermore, there was no significant change in the threat response of animals with constitutive CX3CR1-BAC-Cre expression, as measured by the number of escapes in response to a predator-like looming-shadow stimulus.

#### The constitutive expression of CX3CR1-Cre-recombinase does not have unintended effects in the adult brain, except in the DG of the hippocampus

Early postnatal induction of CX3CR1^CreER(Litt)^ leads to an aberrant microglial phenotype in pups, but this microglial phenotype has been shown to be normalized by adulthood^8^. Thus, we assessed the effects of constitutive CX3CR1-BAC-Cre expression on microglia by immunostaining for IBA1, P2RY12 and CD68 in adult mice (>P60). The goal was to achieve a direct comparison for both microglial markers, IBA1 and P2RY12, in pups and adult mice.

Therefore, Sholl analysis was performed in the set of regions previously mentioned with sex included as a variable. However, P2RY12 staining was observed to be punctate and diffuse in adult animals, unlike in pups, with primary labeling of the processes rather than the soma. Thus, we were unable to perform microglial morphology analysis using P2RY12 in adults.

No changes were found in any parameters for the PC (Fig. 4), PVN (Fig. S12), or CA3 (Fig. S15). The amygdala only exhibited sex differences in the volume of CD68 lysosomes (main effect of Sex, F_(1,28)_=8.64, p<0.05; Fig. S13I), with females being higher overall than males. In a comparable manner, the CA1 region of the hippocampus also only exhibited sex differences.

Females showed an overall increased volume of P2RY12 (main effect of Sex, F_(1,30)_=7.977, p<0.05; Fig. S14H), and decreased volume of CD68 lysosomes (main effect of Sex, F_(1,30)_=7.194, p<0.05; Fig. S14I).

The only region where the effects of constitutive CX3CR1-Cre expression were still detectable was the DG. where Cre+ males had increased IBA1 density in the DG (interaction of Sex x Genotype, F_(1, 29)_=8.964, p<0.05; Šídák, p<0.05; Fig. S16B); as well as increased IBA1 volume (main effect of Genotype, F_(1,30)_=6.128, p<0.05; Fig. S16C). However, no significant changes in morphology or CD68 lysosome volume were found.

## DISCUSSION

The extensive use of inducible and constitutive CX3CR1-Cre models in studying microglia have permitted significant advancement in our understanding of microglial biology; however, off-target effects need to be thoroughly assessed to guarantee the validity of the results. Previous studies have characterized the unintended effects of inducible CX3CR1^CreER^ expression in microglia and have warned against its use, particularly with induction during early postnatal ages due to its long-lasting and detrimental effects on anxiety-like behavior. Here, we studied the possible unintended effects of constitutive CX3CR1-BAC-Cre expression. We hypothesized that due to its uninterrupted and sustained expression throughout an animal’s life, constitutive CX3CR1-BAC-Cre expression would lead to adverse and off-target effects on microglia and adult mouse behavior that are potentially even more detrimental than the inducible Cre model. Thus, we sought to thoroughly characterize the effects of constitutive CX3CR1-BAC-Cre expression on microglia and rigorously compared CX3CR1-BAC-Cre+/- and CX3CR1-BAC-Cre-/-male and female littermates at two time-points: early postnatal age (P8±1) and adulthood (>P60).

During the early postnatal age, we observed a subtle change in microglial phenotype with slightly decreased microglial volume in the PC, PVN, and CA1 of the Hippocampus of CX3CR1-Cre+/-animals. However, alterations in microglial morphology alone cannot predict a change in function^2^. The changes in volume or density were different depending on the microglial marker (IBA1 or P2RY12); this is expected, as different subsets of microglia can express each marker in distinct levels. It is known that, especially during the early postnatal period, microglia are highly heterogeneous throughout the brain^16^. Therefore, this result might point out that subgroups of microglia are affected differently by Cre expression.

Beyond subpopulations, we demonstrated that regional differences in the context of microglia are especially important. We found that the hippocampus was particularly vulnerable to the expression of Cre, since it caused a decrease in the engulfment of PSD95 and gephyrin puncta in the CA1, as well as a decrease in the volume of IBA1 in males in the DG during adulthood.

This might be related to the high plasticity and neurogenesis that characterizes this region. Since the hippocampus recapitulates the developmental trajectory throughout life, it is thought to be particularly vulnerable to insults^17^. Interestingly, the aberrant early postnatal microglial volume in females in the CA1 was normalized by adulthood, which could also indicate a transient shift in the developmental trajectory. Conversely, the IBA1 volume was only affected in adulthood in the DG, which might suggest that microglia from this region have a later sensitive period to Cre, beyond our developmental timepoint (P8±1). Notably, unlike inducible Cre, constitutive CX3CR1-Cre expression did not lead to any changes in adult mouse behavior, as tested with the open-field test, elevated-plus maze test, and looming-shadow threat task.

Overall, our results suggest that the effects of constitutive CX3CR1-BAC-Cre expression on microglia are far less noticeable than inducible CX3CR1-Cre expression, although effects can vary by age, brain region, and sex, which aligns well with the well-known heterogeneity of microglial populations^16^.

We propose that the difference in the genetic targeting strategy of these transgenic lines may explain the observed differences in their off-target effects. The constitutive CX3CR1-Cre transgenic line used for this study employs BAC genetic engineering, which inserts the transgene Cre cassette via random integration into the genome and maintains diploidy. To avoid an unwanted phenotype from the random integration, the constitutive CX3CR1-BAC-Cre mouse line is utilized in its heterozygous state. In contrast, the inducible CX3CR1-Cre (CX3CR1^CreER(Litt)^) employs the “knock-in” strategy. Thus, even when bred to heterozygosity, CX3CR1^CreER^ disrupts the diploidy of CX3CR1 endogenous gene and inserts the conditionally active Cre cassette into the CX3CR1 locus. The CX3CR1 receptor is involved in neuron-microglia fractalkine signaling, which plays a vital role in microglial developmental synaptic pruning and homeostasis. Its loss and deficiency can alter microglial number, subsequently altering microglia-mediated synaptic pruning during developmental ages^13,18^. Therefore, it is possible that the heterozygous deletion of CX3CR1-Cre in the inducible model could have produced more highly reactive microglia in specific brain regions.

Our study demonstrates that constitutive CX3CR1-BAC-Cre expression leads to subtle alterations in microglial phenotype in early postnatal ages, with no alterations in adult mouse behavior and enduring microglial changes only in the hippocampus. This finding contrasts with the key finding from Sahasrabuddhe et al.^8^, which shows that early postnatal induction of CX3CR1^CreER^ leads to IFN-1 signaling activation, increased developmental synaptic pruning in the somatosensory cortex, and increased anxiety-like behavior in adulthood. A thorough comparison of Cre-recombinase levels between both lines across different time points is required to conclude direct causal effects. We predict that a sudden burst in Cre-recombinase levels with exogenous induction, especially during early postnatal ages, might alert microglia to react to increased levels as if it were an immune challenge, further impairing their function during critical development. In contrast, constitutive presence and gradual accumulation of Cre-recombinase over an animal’s life can result in microglia adapting to these signals. Notably, unlike CX3CR1^CreER(Litt)^, CX3CR1^CreER(Jung)^ has been shown to have minimal off-target effects, with no alterations in homeostatic markers of microglia following early postnatal induction^9^. As proposed earlier, the genetic targeting strategy can help explain the observed differences; CX3CR1^CreER(Litt)^ expresses a Cre fusion protein along with enhanced yellow fluorescent protein (EYFP), making it a larger protein construct, which might make it more likely to trigger an immune response.

The Cre-loxP system has proven to be an indispensable genetic tool due to the spatial and temporal specificity it provides. Nevertheless, our findings and previous studies^8,9^ recognize the drawbacks of adverse and off-target effects, especially when employing this tool to study naturally reactive cells, such as microglia. While these studies encourage new advances in perfecting Cre-loxP genetic technology or alternatives, it becomes more crucial than ever to include correct controls in experimental studies as we continue to use the existing Cre transgenic lines. We recommend breeding these lines to heterozygosity and running littermate control validation experiments (Cre+/-vs Cre-/-) even before crossing with loxP lines, to determine if there is a baseline alteration caused by the expression of Cre alone.

### Limitations of the study

In this study, we did not test for direct effects of Cre expression on the immune function of microglia. A detailed analysis of the effects of constitutive CX3CR1-BAC-Cre on microglia-mediated IFN-1 signaling activation may provide further insights, although we did not detect significant DNA damage in our model, which was the trigger for the IFN-1 signaling in the inducible Cre model^8^. The phosphorylated-gamma-H2AX antibody used to detect DNA damage here was the same as used in the previous study^8^, but we found that it generally produced diffuse and non-optimal staining results. Future studies could alternatively employ TUNEL staining to confirm that DNA damage is not induced by constitutive CX3CR1-Cre expression.

## ACKNOWLEDGMENTS

This work was supported by NIH grant R00 MH120327 (JLB) and the Whitehall Foundation. Research reported in this publication was supported by the Office of The Director, National Institutes Of Health and National Institute Of General Medical Sciences under Award Number S10OD032336-01. The content is solely the responsibility of the authors and does not necessarily represent the official views of the National Institutes of Health. We thank Logan Ouellette and Sylvie Call for excellent technical assistance, and the Georgia State University Division of Animal Resources for outstanding animal care. We also acknowledge Hannah Lichtenstein for feedback on the results and assistance with editing the manuscript.

## AUTHOR CONTRIBUTIONS

FHMR, BD, JS, and JLB designed the experiments. FHMR, BD, MG, JS, FK, and UK performed the experiments. FHMR, BD, MG, JS, FK, UK, and JLB analyzed data. FHMR, BD, MG, and JLB wrote and/or edited the paper.

## METHODS

### Experimental Model and Study Participants Details Animals

Both males and females of the CX3CR1-BAC-Cre ([Tg (Cx3cr1-Cre) MW126Gsat], GENSAT Transgenic Project) transgenic mouse line were used for all experiments with the Cre-recombinase enzyme expressed constitutively under the direction of the CX3CR1 promoter. CX3CR1-BAC-Cre+/-male mice were bred with CX3CR1-BAC-Cre-/-female mice to obtain CX3CR1-BAC-Cre-/- and CX3CR1-BAC-Cre+/-experimental animals. Mice were housed under 12 h light/dark cycle with free access to food and water. All experiments were performed in accordance with National Institutes of Health (NIH) guidelines and were approved by the Georgia State University Animal Care and Use Committee.

### Experimental Design

The effects of constitutive CX3CR1-BAC-Cre expression were analyzed in CX3CR1-BAC-Cre-/- and CX3CR1-BAC-Cre+/-pups (P8±1 d) and adults (>P60). The pups (P8±1 d) were euthanized with Euthasol (Patterson Veterinary, VIRBAC Animal Health) and perfused transcardially with ice-cold 1x phosphate-buffer saline (PBS) followed by 4% paraformaldehyde. Perfused brains were then post-fixed in 4% paraformaldehyde for 4 hours, then dehydrated in 0.1 M PBS 15% sucrose solution overnight followed by 25% (30% for adults) sucrose solution until brains sank. Brains were frozen by dipping into 2-methylbutane on dry ice for ∼60 s and then stored in a -80°C freezer. The cohort of adults (>P60) was subjected to a series of behavioral assays: looming-shadow threat task, open field test, elevated-plus maze, and then perfused as described above.

### Method Details Immunohistochemistry

Pup and adult brains were coronally sectioned into 25-μm-thick slices using a Leica cryostat and stored in an anti-freeze solution in a -20°C freezer. For immunohistochemistry, floating brain sections were washed several times (3x, five minutes each) with PBS-T (PBS containing 0.3% Triton X-100; pH=7.4) at room temperature (RT). Tissues were then allowed to permeabilize in 0.09% H_2_O_2_ in PBS-T for 20 min at RT. The sections were then incubated in blocking solution 5% normal donkey serum (NDS) for 1 h at RT and then stained in primary antibody solution (in 2% NDS) overnight at 4°C. Sections were then washed in PBS-T several times and incubated in secondary antibody solution (in 2% NDS) for 3 h at RT. After repeated PBS-T washes, free floating sections were counter-stained with DAPI for 1 min before mounting on gelatin-coated slides and cover-slipping. Immunostaining P8 brain sections with chicken anti-IBA1 required antigen-retrieval with Tris-EDTA for 3 mins at 90°C before the blocking step.

**Table 1.** Antibodies used for immunohistochemistry.

| Catalog number | Primary antibody | Catalog number | Corresponding secondary antibody |
| --- | --- | --- | --- |
| AS-55043A | Rabbit anti-P2RY12 (1:2000) | NC0254454 | Donkey anti-rabbit Alexa Fluor 647 (1:1000) |
| 50-164-936 | Rat anti-CD68 (1:2000) | A21208 | Donkey anti-rat Alexa Fluor 488 (1:500) |
| 234 009 | Chicken anti-IBA1 (1:5000) | 703-605-155 | Donkey anti-chicken Alexa Fluor 647 (1:1000) |
| 14569882 | Rat anti-Ki67 (1:1000) | A21208 | Donkey anti-rat Alexa Fluor 488 (1:500) |
| SAB5700329 | Rabbit anti-phosphorylated-Gamma-H2AX (1:500) | A10042 | Donkey anti-rabbit Alexa Fluor 568 (1:500) |
| 234 009 | Chicken anti-IBA1(1:5000) | 703-605-155 | Donkey anti-chicken Alexa Fluor 647 (1:1000) |
| NC0292731 | Guinea pig anti-Gephyrin (1:500) | 706-545-148 | Donkey anti-guinea pig Alexa Fluor 488 (1:500) |
| 51-690-0 | Rabbit anti-PSD95 (1:1000) | A10042 | Donkey anti-rabbit Alexa Fluor 568 (1:1000) |
| 50-164-936 | Rat anti-CD68 (1:2000) | 712-605-153 | Donkey anti-rat Alexa Fluor 647 (1:1000) |
| 50-164-936 | Rat anti-CD68 (1:2000) | A21208 | Donkey anti-rat Alexa Fluor 488 (1:500) |
| 234 009 | Chicken anti-IBA1(1:2000) | A78950 | Donkey anti-chicken Alexa Fluor 568 (1:1000) |
| AS-55043A | Rabbit anti-P2RY12 (1:2000) | NC0254454 | Donkey anti-rabbit Alexa Fluor 647 (1:1000) |

### Confocal Imaging and Analysis

Confocal images of the canonical microglial markers during early postnatal development were acquired from the parietal cortex (PC), the paraventricular nucleus of the hypothalamus (PVN), the amygdala, and hippocampal subregions (CA1, CA3, Dentate gyrus) using a Zeiss LSM-700 confocal microscope. Each image consisted of a z-stack of 13 planes with an interval of 1μm using the 20X objective. The rest of the images (microglial proliferation and DNA damage assessment, synaptic puncta, and microglial markers in adulthood) were obtained using an LSM-780 confocal microscope. The same objective and z-stack configuration was used, with the exception of synaptic puncta imaging, which required a 63X objective and intervals of 0.5 μm. FIJI software was used to manually count IBA1-positive and P2RY12-positive microglia, Ki67-positive proliferating cells and phosphorylated-Gamma-H2AX-positive microglia. Microglia volume and morphology were analyzed using 3-D reconstruction and Filament tool in Bitplane Imaris software version 9.3.1 for every brain region except PC, where microglial morphology was reconstructed using the filament tracing tool from version 10.2.0 since the other brain regions were analyzed before the launch if version 10.2.0. This newer version of Imaris allowed us to obtain finer details from the Sholl analysis such as soma vs. segment volume; additionally, it incorporated machine learning for proper detection of microglia. For each animal, three bilateral (left and right) technical replicates were analyzed in the PC, and two for every other brain region. The average value of the technical replicates was used to represent that animal in each variable. Measures of Sholl intersections, total branch points and maximum filament length from Sholl analysis^19^ were acquired using the Imaris Filament tool with 1-μm concentric circles placed around IBA1-positive and P2RY12-positive microglia. Imaris’ Surface tool was used to measure the count and volume of CD68-positive lysosomes. Imaris’ Spots tool was used to count the PSD95-positive and Gephyrin-positive excitatory and inhibitory post-synaptic structures, respectively. Additionally, object-object statistics from the post-synaptic markers to CD68 surfaces were used in the PC to quantify microglial post-synaptic engulfment.

### Behavioral Assays

One week prior to behavioral testing, mice were transferred to a 12 h reverse light cycle room. Each task was conducted in a quiet room, illuminated only with red light where subjects were allowed 1 hour to acclimate in their home cage before any behavioral manipulation. All testing occurred in adulthood (>P60), with male and female mice being tested on separate days for each task.

### Looming-Shadow Threat Task

The looming-shadow threat task assesses a mouse’s response to a predator-like threat^20^. Prior to testing, mice were habituated to an arena (43cm x 18cm x 16 cm) containing a plastic shelter, painted matte black, for 15 min. Following habituation, a looming-shadow stimulus was presented five times to the mouse from above with at least 1 min and 15 s in between each stimulus. The looming stimulus consisted of a small black circle displayed on a monitor placed on the top of the arena that was stationary for 3 s, expanded for 2 s and then stationary for another 3 s. Responses to the stimulus were scored as escaping, freezing, or no response and later verified on a video recording. An additional 6 s following the stimulus presentation was scored due to the observation of delayed escape behavior. The following criteria were used to differentiate between the behaviors: 1) any flight response resulting in the mouse reaching the shelter within 14 s of the start of the stimulus was classified as escape; 2) all cessation of movement aside from breathing was classified as freezing; 3) the presentation of any behavior that did not classify as escaping or freezing within 14 s of the start of the stimulus was classified as no response. Escape behavior is considered to be an active defensive response dependent on increased PVN-CRH+ neuron activity^20^. Percentage of each response type as well as latency to respond to the looming stimulus was analyzed for each subject. If no response was observed, latency was recorded as 14 s.

### Open Field Test

One week following the looming-shadow threat task, mice underwent a 10-minute open field test (OFT). The arena consisted of a 43 cm × 43 cm floor surrounded by a 30 cm wall that was encased in a wooden box with double doors. Med Associates software was used to divide the arena into 225 equivalent squares with the 144 outermost squares classified as the “surround” zone and the 81 innermost squares classified as the “center” zone. Total distance travelled as well as the percentage of time subjects spent in the surround and the center zone was analyzed. Reduced time spent in the center zone is considered to be a marker of anxiety-like behavior^8,21^.

### Elevated-Plus Maze Test

Three days following the OFT, mice underwent a 5 min elevated-plus maze test. The apparatus was comprised of two closed arms (35 × 5 × 16.5 cm) and two open arms (35 × 5 × 2.5 cm) intersecting perpendicularly and elevated 60 cm off the floor. Mice were placed into the center of the arena with their head positioned towards a closed arm at the start of each test. ANY-maze software (Stoelting) was used to track the center of each subject as it moved freely throughout the apparatus. Recordings were analyzed for total distance travelled and percentage of time spent in each arm and the intersecting center. Reduced time spent exploring the open arms is used as a marker for anxiety-like behavior^8,22^.

### Quantification and Statistical Analysis

Statistical differences between CX3CR1-BAC-Cre-/- and CX3CR1-BAC-Cre+/-groups were assessed using GraphPad Prism software (GraphPad, San Diego, CA, USA). Each group consisted of n=7-11 animals. 2-way ANOVA tests were used to analyze the effects of constitutive CX3CR1-BAC-Cre using genotype (Cre-/- and Cre+/-) and sex as independent variables. The dependent variables were microglial volume, density, proliferation, DNA damage, and post-synaptic structures numbers. 3-way repeated-measures ANOVA tests were used to analyze the effects of constitutive CX3CR1-BAC-Cre expression and sex on microglial morphology via Sholl intersections at 1-μm radius intervals for IBA1-positive and P2RY12-positive microglial processes. If found significant, the analysis of Sholl Intersections was further split by sex. Grubbs’ test was used to remove outliers. Significant interactions were followed by Šídák post hoc tests. The significance level was set to 0.05 for all tests, and data are presented as mean ± SEM. All experiments were assessed blindly without prior knowledge of the experimental group.

