## Supplemental Information for "Constitutive expression of CX3CR1-BAC-Cre introduces minimal off-target effects in microglia"

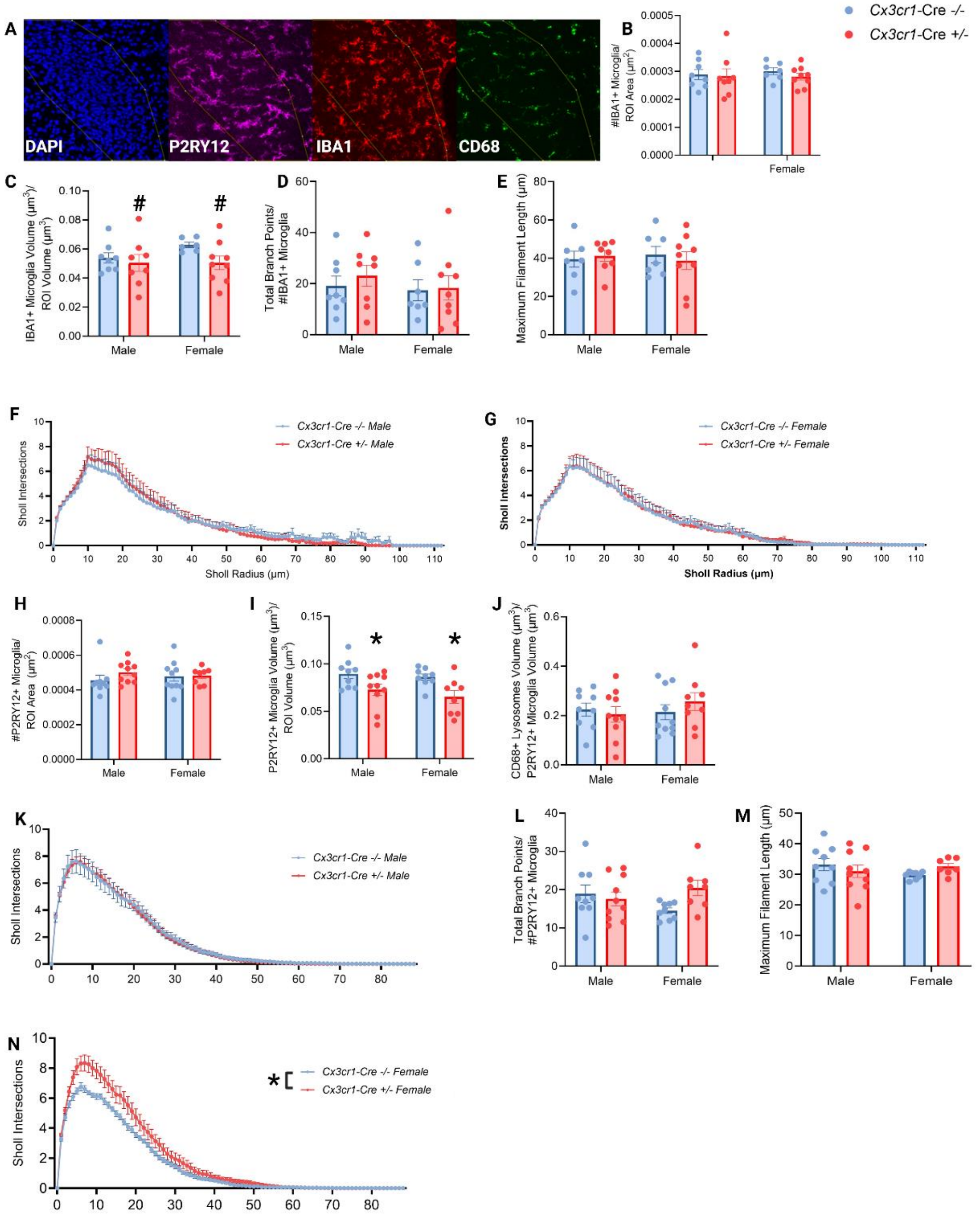


**Figure S1. Effects of constitutive CX3CR1-BAC-Cre expression on early postnatal microglia in the PVN.**

- **A.** Representative images showing DAPI, P2RY12, IBA1 and CD68 staining in the PVN.
- **B.** Quantification of IBA1+ microglia density.
- **C.** Quantification of IBA1+ microglia volume (trend for main effect of genotype, $F_{(1,27)}$=2.945, p=0.098)

**D.** (C-F) IBA1 Sholl analysis metrics. Total branch points of IBA1+ microglia.
**E.** Maximum filament length of IBA1+ microglia.
**F.** Number of Sholl intersections for each radius of IBA1+ microglia of male mice.
**G.** Number of Sholl intersections for each radius of IBA1+ microglia of female mice.
**H.** Quantification of P2RY12+ microglia density.
**I.** Quantification of P2RY12+ microglia volume (main effect of genotype, $F_{(1,16)}=6.886$, $p<0.05$)
**J.** Quantification of CD68+ microglial lysosomes volume relative to the P2RY12+ microglial volume.
**K.** (J-M) P2RY12 Sholl analysis metrics. Number of Sholl intersections for each radius of P2RY12+ microglia of male mice.
**L.** Total branch points of P2RY12+ microglia.
**M.** Maximum filament length of P2RY12+ microglia.
**N.** Number of Sholl intersections for each radius of P2RY12+ microglia of female mice (main effect of genotype, $F_{(1,16)}=6.886$, $p<0.05$)
Each dot represents the average value for one animal. n=7-11. *p<0.05. Mean ± SEM.

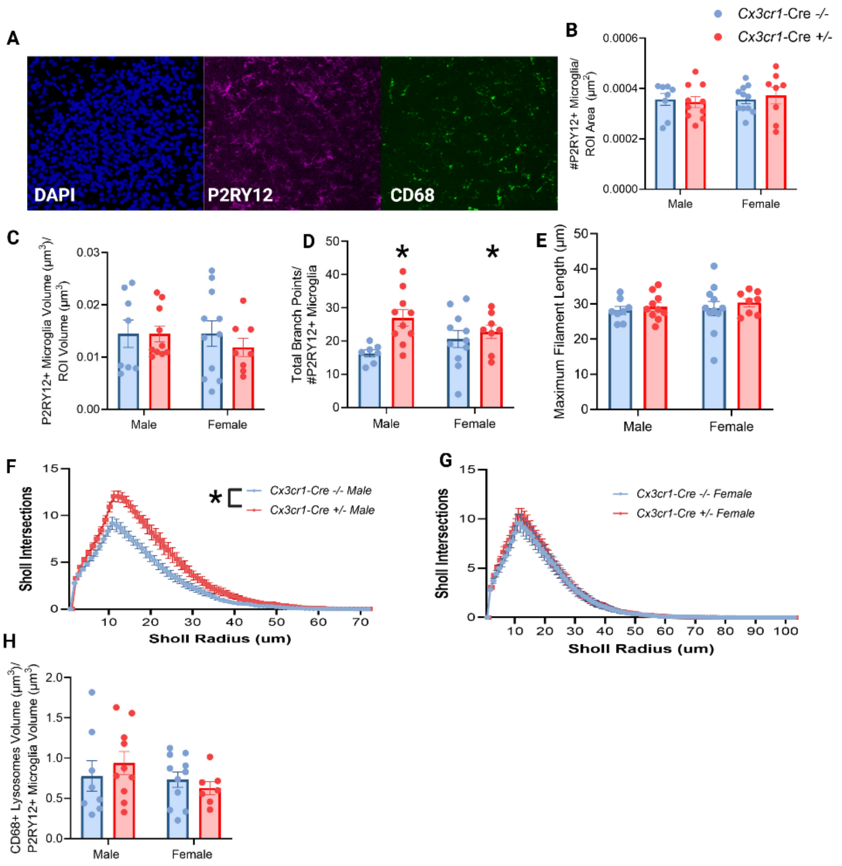


**Figure S2. Effects of constitutive CX3CR1-BAC-Cre expression on early postnatal microglia in the Amygdala.**

- **A.** Representative images showing DAPI, P2RY12, and CD68 staining in the Amygdala.
- **B.** Quantification of P2RY12+ microglia density.
- **C.** Quantification of P2RY12+ microglia volume.
- **D.** (C-F) P2RY12 Sholl analysis metrics. Total branch points of P2RY12+ microglia.
- **E.** Maximum filament length of P2RY12+ microglia.
- **F.** Number of Sholl intersections for each radius of P2RY12+ microglia of male mice (interaction of Distance from soma x Genotype x Sex, $F_{(102,3399)}=1.532$, $p<0.05$; 2-way ANOVA, $F_{(1,16)}=9.000$, $p<0.05$)
- **G.** Number of Sholl intersections for each radius of P2RY12+ microglia of female mice
- **H.** Quantification of CD68+ microglial lysosomes volume relative to the P2RY12+ microglial volume.

  Each dot represents the average value for one animal. n=7-11. *p<0.05. Mean ± SEM.

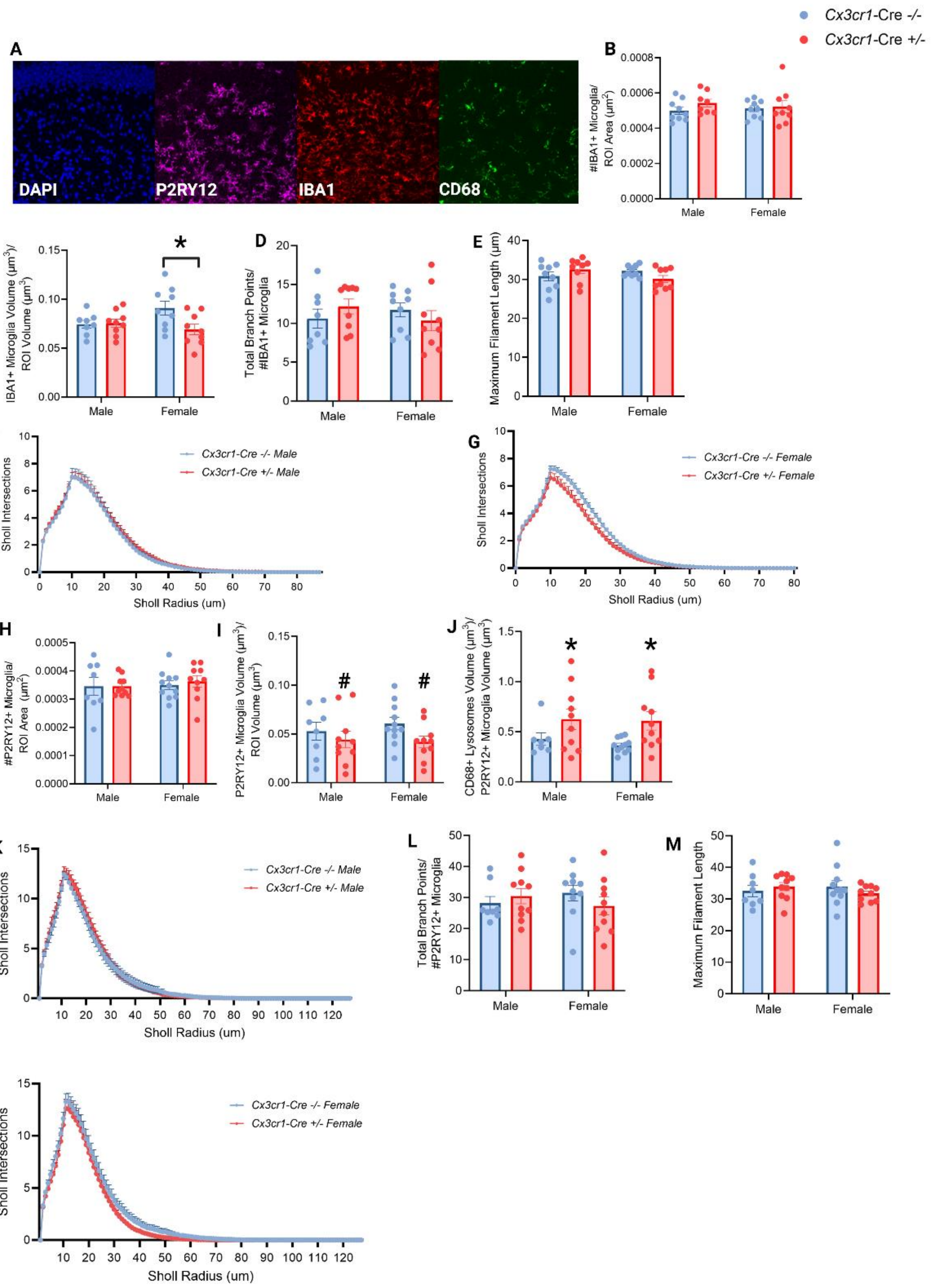


**Figure S3. Effects of constitutive CX3CR1-BAC-Cre expression on early postnatal microglia in the CA1.**

- **A.** Representative images showing DAPI, P2RY12, IBA1 and CD68 staining in the CA1.
- **B.** Quantification of IBA1+ microglia density.
- **C.** Quantification of IBA1+ microglia volume (interaction of Genotype x Sex $F_{(1,31)}$=4.50, $p<0.05$, Šídák, $p<0.05$)
- **D.** (C-F) IBA1 Sholl analysis metrics. Total branch points of IBA1+ microglia.

**E.** Maximum filament length of IBA1+ microglia.
**F.** Number of Sholl intersections for each radius of IBA1+ microglia of male mice.
**G.** Number of Sholl intersections for each radius of IBA1+ microglia of female mice.
**H.** Quantification of P2RY12+ microglia density.
**I.** Quantification of P2RY12+ microglia volume (trend for main effect of genotype, $F_{(1,35)}$=3.33, p=0.07)
**J.** Quantification of CD68+ microglial lysosomes volume relative to the P2RY12+ microglial volume (main effect of Genotype $F_{(1,34)}$=7.986; p<0.05)
**K.** (J-M) P2RY12 Sholl analysis metrics. Number of Sholl intersections for each radius of P2RY12+ microglia of male mice.
**L.** Total branch points of P2RY12+ microglia.
**M.** Maximum filament length of P2RY12+ microglia.
**N.** Number of Sholl intersections for each radius of P2RY12+ microglia of female.
Each dot represents the average value for one animal. n=7-11. *p<0.05. Mean ± SEM.

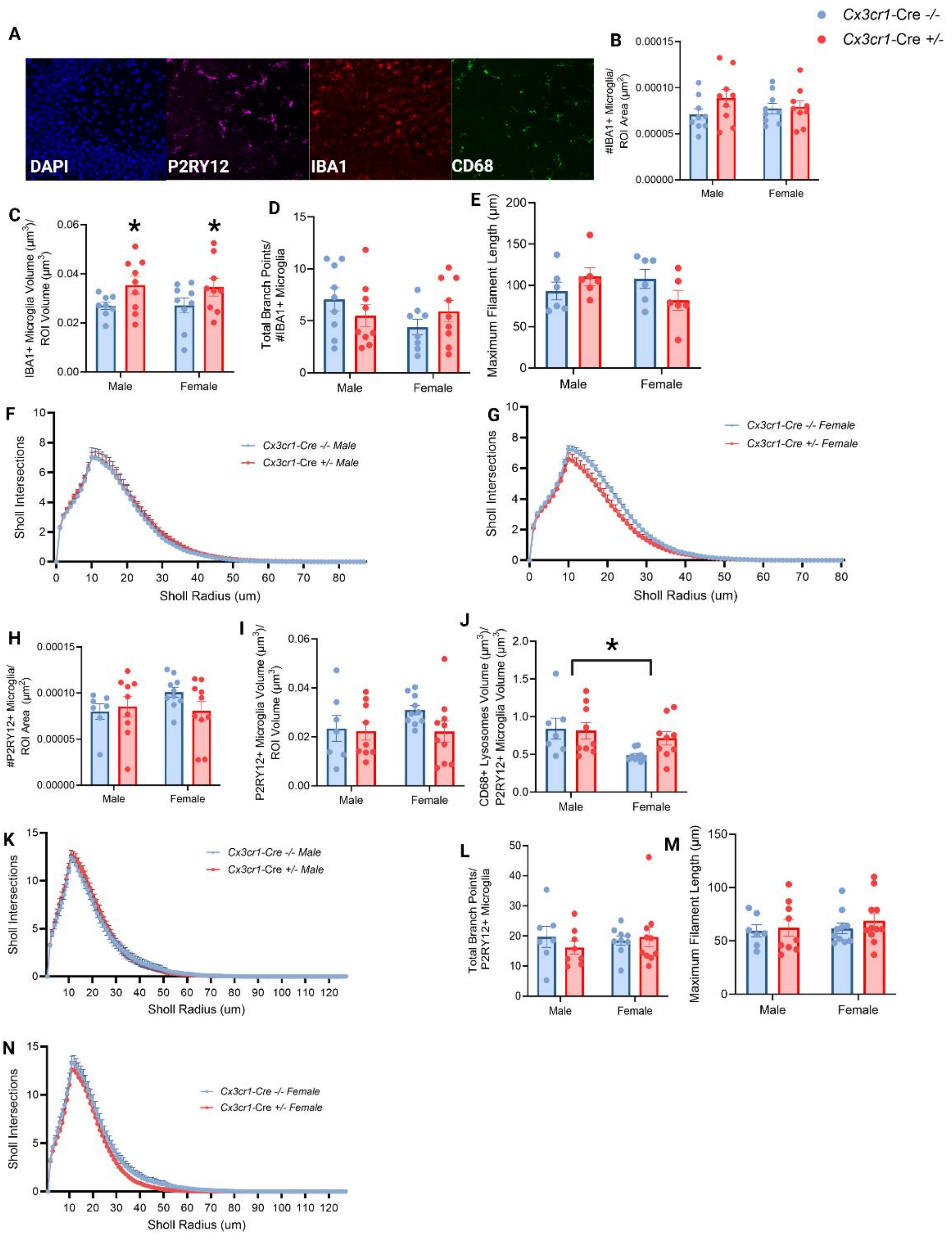


**Figure S4. Effects of constitutive CX3CR1-BAC-Cre expression on early postnatal microglia in the CA3.**

- **A.** Representative images showing DAPI, P2RY12, IBA1 and CD68 staining in the CA3.
- **B.** Quantification of IBA1+ microglia density.
- **C.** Quantification of IBA1+ microglia volume (main effect of Genotype, $F_{(1,31)}=6.28$, $p<0.05$)
- **D.** (D-G) IBA1 Sholl analysis metrics. Total branch points of IBA1+ microglia.

**E.** Maximum filament length of IBA1+ microglia.
**F.** Number of Sholl intersections for each radius of IBA1+ microglia of male mice.
**G.** Number of Sholl intersections for each radius of IBA1+ microglia of female mice.
**H.** (G-I) P2RY12 metrics. Quantification of P2RY12+ microglia density.
**I.** Quantification of P2RY12+ microglia volume.
**J.** Quantification of CD68+ microglial lysosomes volume relative to the P2RY12+ microglial volume (main effect of Sex $F_{(1,30)}=5.752$; $p<0.05$)
**K.** (K-N) P2RY12 Sholl analysis metrics. Number of Sholl intersections for each radius of P2RY12+ microglia of male mice.
**L.** Total branch points of P2RY12+ microglia.
**M.** Maximum filament length of P2RY12+ microglia.
**N.** Number of Sholl intersections for each radius of P2RY12+ microglia of female.
Each dot represents the average value for one animal. n=7-11. *$p<0.05$. Mean ± SEM.

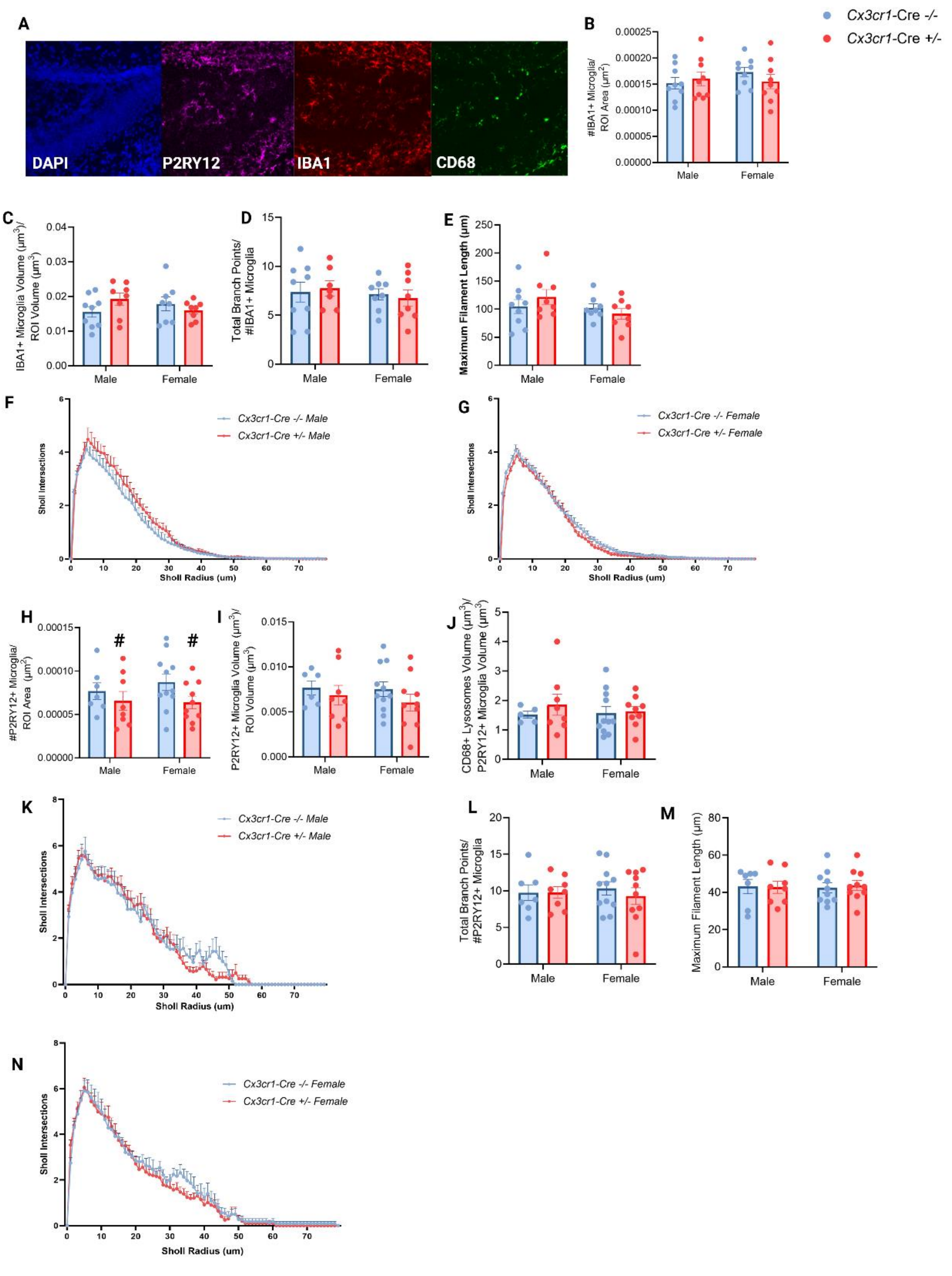


**Figure S5. Effects of constitutive CX3CR1-BAC-Cre expression on early postnatal microglia in the DG.**

- **A.** Representative images showing DAPI, P2RY12, IBA1 and CD68 staining in the DG.
- **B.** Quantification of IBA1+ microglia density.
- **C.** Quantification of IBA1+ microglia volume.
- **D.** (D-G) IBA1 Sholl analysis metrics. Total branch points of IBA1+ microglia.

**E.** Maximum filament length of IBA1+ microglia.

**F.** Number of Sholl intersections for each radius of IBA1+ microglia of male mice.

**G.** Number of Sholl intersections for each radius of IBA1+ microglia of female mice.

**H.** P2RY12 metrics. Quantification of P2RY12+ microglia density (trend for main effect of genotype, $F_{(1,32)}$=3.260, p=0.0804).

**I.** Quantification of P2RY12+ microglia volume.

**J.** Quantification of CD68+ microglial lysosomes volume relative to the P2RY12+ microglial volume.

**K.** (K-N) P2RY12 Sholl analysis metrics. Number of Sholl intersections for each radius of P2RY12+ microglia of male mice.

**L.** Total branch points of P2RY12+ microglia.

**M.** Maximum filament length of P2RY12+ microglia.

**N.** Number of Sholl intersections for each radius of P2RY12+ microglia of female.
Each dot represents the average value for one animal. n=7-11. *p<0.05. Mean ± SEM.

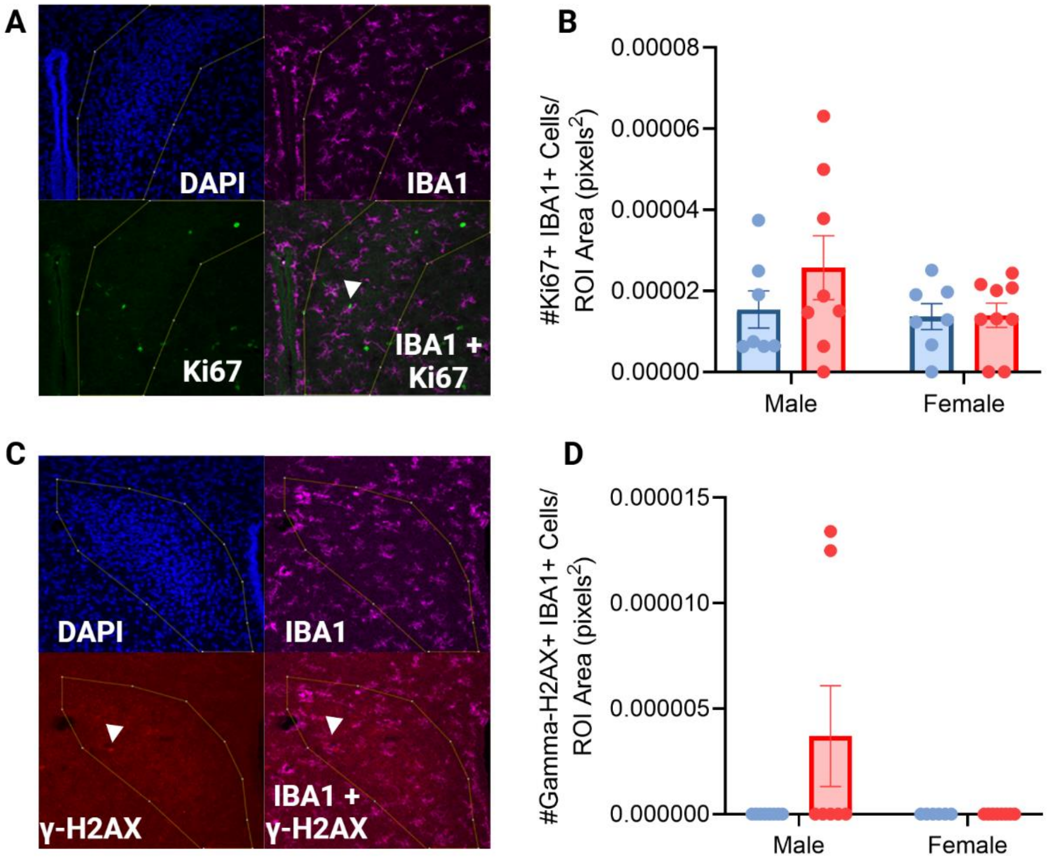


**Figure S6. Effects of constitutive CX3CR1-BAC-Cre expression on proliferation and DNA damage of early postnatal microglia in the PVN.**

A. Representative images showing PVN stained with DAPI, Ki67 and IBA1. White arrowhead points to a cell double-positive for Ki67 and IBA1.
B. Quantification of density of cells double-positive for Ki67 and IBA1.
C. Representative image showing a PVN stained with DAPI, IBA1 and phosphorylated-γ-H2AX. White arrowhead points to a cell double-positive for IBA1 and phosphorylated-γ-H2AX.
D. Quantification of density of cells double-positive for phosphorylated-γ-H2AX and IBA1. Each dot represents the average value for one animal. n=7-9. Mean ± SEM.

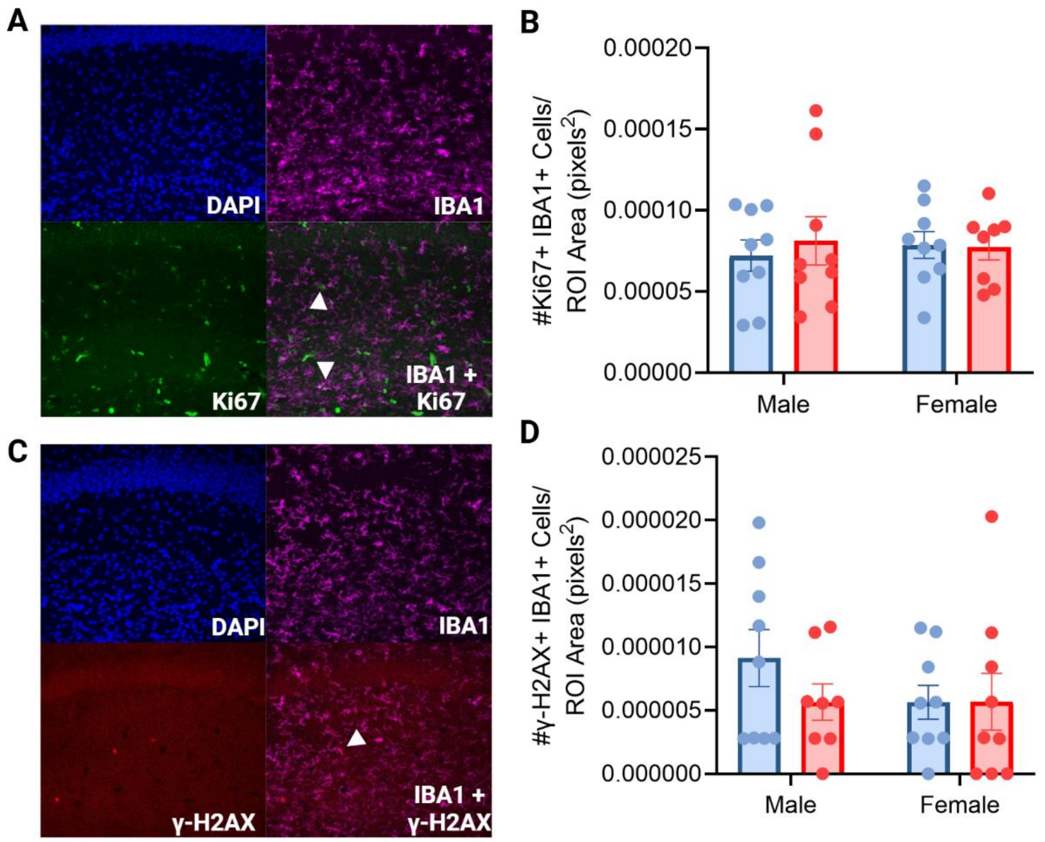


**Figure S7. Effects of constitutive CX3CR1-BAC-Cre expression on proliferation and DNA damage of early postnatal microglia in the CA1.**

A. Representative images showing the CA1 stained with DAPI, Ki67 and IBA1. White arrowhead points to a cell double-positive for Ki67 and IBA1
B. Quantification of density of cells double-positive for Ki67 and IBA1.
C. Representative image showing the CA1 stained with DAPI, IBA1 and phosphorylated-γ-H2AX. White arrowhead points to a cell double-positive for Ki67 and phosphorylated-γ-H2AX
D. Quantification of density of cells double-positive for phosphorylated-γ-H2AX and IBA1. Each dot represents the average value for one animal. n=7-9. Mean ± SEM.

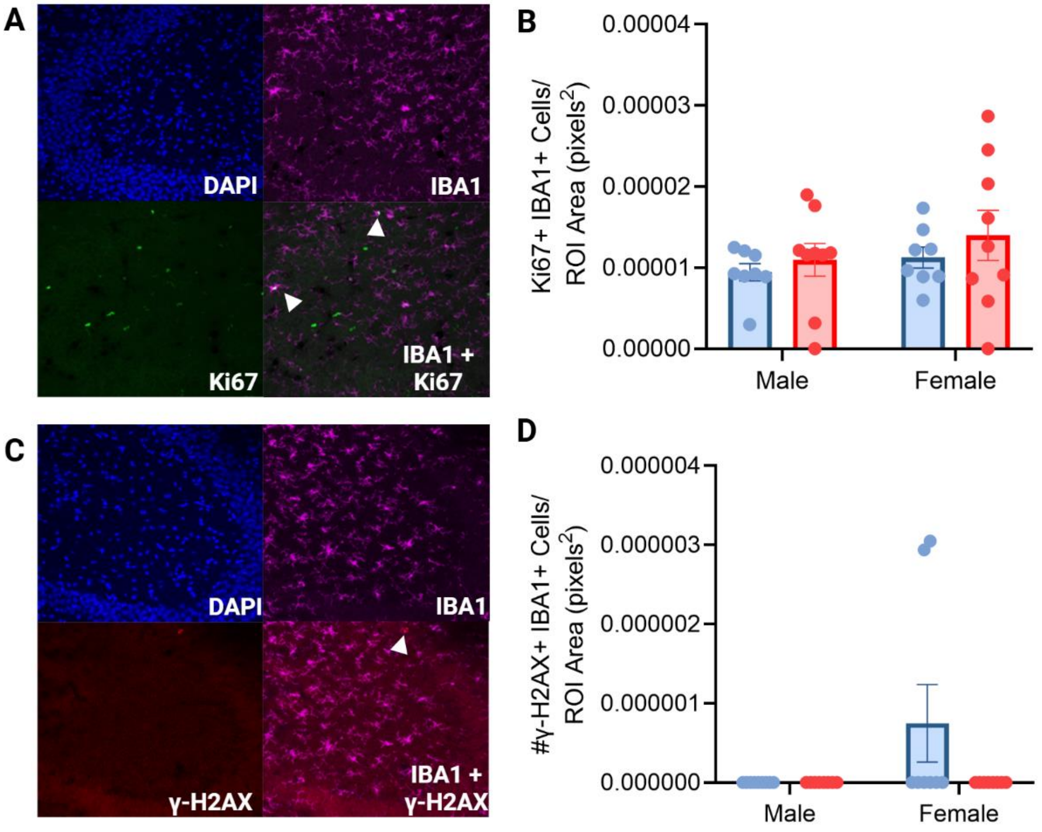


**Figure S8. Effects of constitutive CX3CR1-BAC-Cre expression on proliferation and DNA damage of early postnatal microglia in the CA3.**

A. Representative images showing the CA3 stained with DAPI, Ki67 and IBA1. White arrowhead points to a cell double-positive for Ki67 and IBA1.
B. Quantification of density of cells double-positive for Ki67 and IBA1.
C. Representative image showing the CA3 stained with DAPI, IBA1 and phosphorylated-γ-H2AX. White arrowhead points to a cell double-positive for Ki67 and phosphorylated-γ-H2AX.
D. Quantification of density of cells double-positive for phosphorylated-γ-H2AX and IBA1. Each dot represents the average value for one animal. n=7-9. Mean ± SEM.

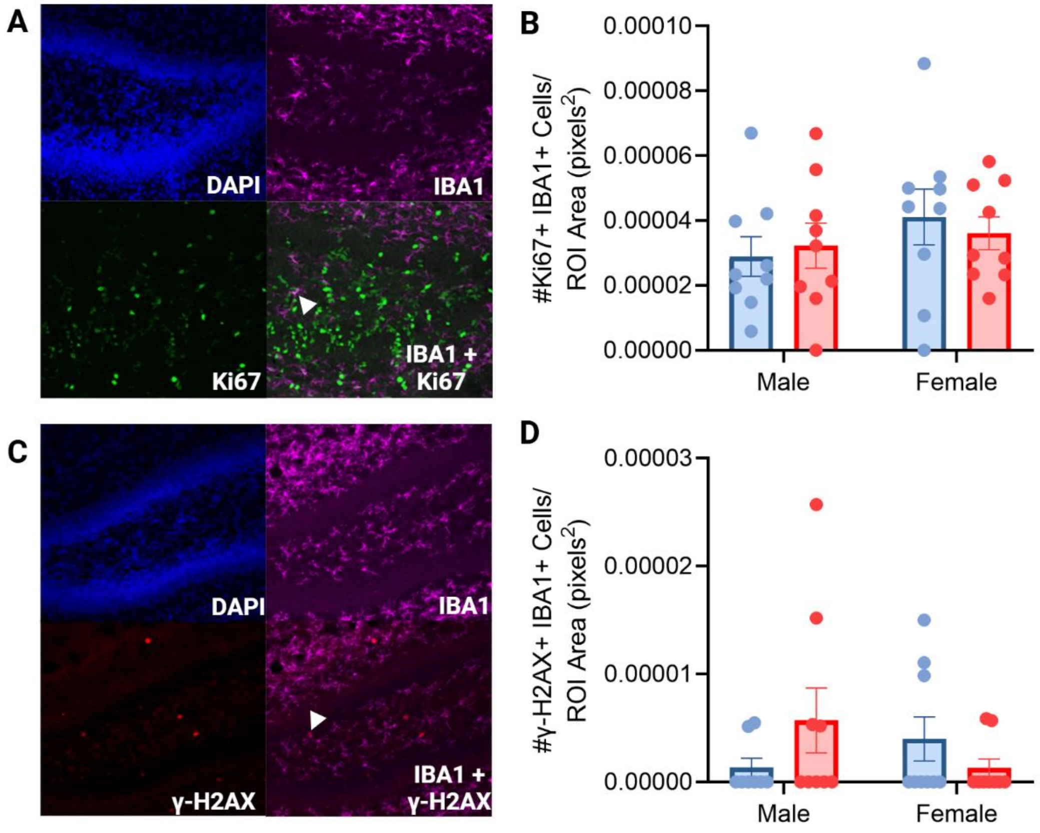


**Figure S9. Effects of constitutive CX3CR1-BAC-Cre expression on proliferation and DNA damage of early postnatal microglia in the DG.**

A. Representative images showing the DG stained with DAPI, Ki67 and IBA1. White arrowhead points to a cell double-positive for Ki67 and IBA1.
B. Quantification of density of cells double-positive for Ki67 and IBA1.
C. Representative image showing the DG stained with DAPI, IBA1 and phosphorylated-γ-H2AX. White arrowhead points to a cell double-positive for Ki67 and phosphorylated-γ-H2AX.
D. Quantification of density of cells double-positive for phosphorylated-γ-H2AX and IBA1. Each dot represents the average value for one animal. n=7-9. Mean ± SEM.

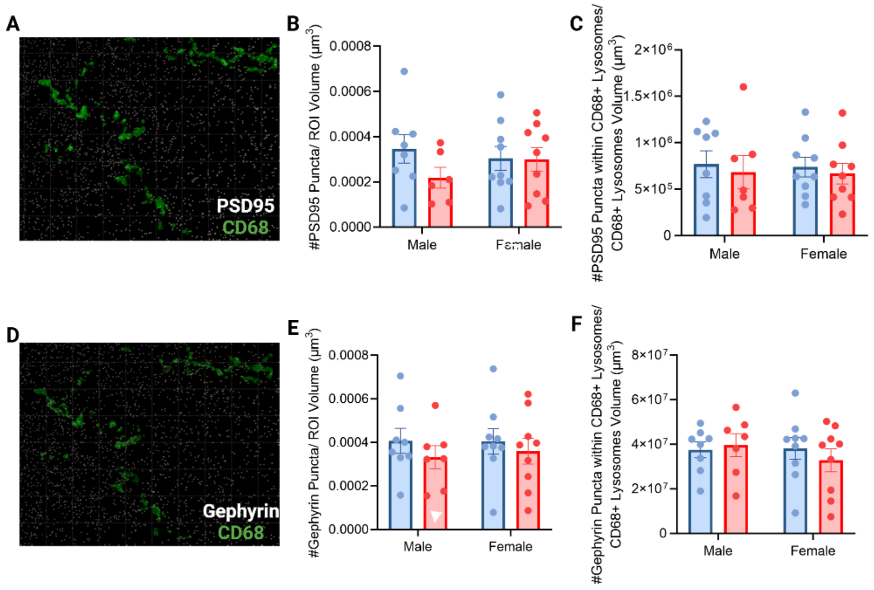


**Figure S10. Effects of constitutive CX3CR1-BAC-Cre expression on synapse engulfment by early postnatal microglia in the PVN.**

A. Representative image showing PSD95 puncta, and the reconstruction of CD68 volume in green. Non-colocalized PSD95 puncta in red, and colocalized puncta in white.
B. Density quantification of PSD95 puncta.
C. Density quantification of PSD95 puncta within CD68+ microglial lysosomes.
D. Representative image showing gephyrin puncta, and the reconstruction of CD68 volume in green. Non-colocalized gephyrin puncta in red, and colocalized puncta in white.
E. Density quantification of gephyrin puncta.
F. Density quantification of gephyrin puncta within CD68+ microglial lysosomes.
Each dot represents the average value for one animal. n=7-9. Mean ± SEM.

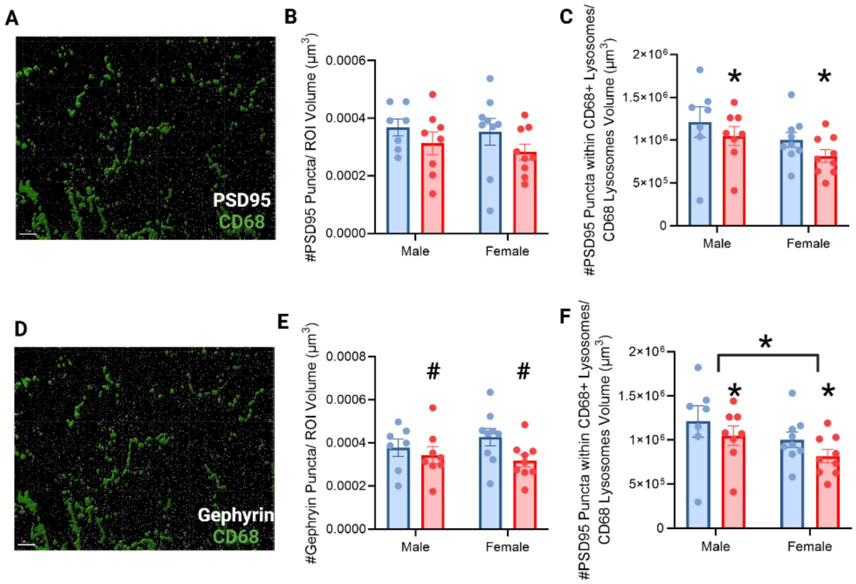


**Figure S11. Effects of constitutive CX3CR1-BAC-Cre expression on synapse engulfment by early postnatal microglia in the CA1.**

A. Representative image showing PSD95 puncta, and the reconstruction of CD68 volume in green. Non-colocalized PSD95 puncta in red, and colocalized puncta in white.
B. Density quantification of PSD95 puncta.
C. Density quantification of PSD95 puncta within CD68+ microglial lysosomes (main effect of genotype, $F_{(1,29)}$=4.453, p<0.05)
D. Representative image showing gephyrin puncta, and the reconstruction of CD68 volume in green. Non-colocalized gephyrin puncta in red, and colocalized puncta in white.
E. Density quantification of gephyrin puncta (trend for main effect of genotype, $F_{(1,29)}$=3.691, p=0.0646)
F. Density quantification of gephyrin puncta within CD68+ microglial lysosomes (main effect of sex, $F_{(1,29)}$=6.245, p<0.05; main effect of genotype, $F_{(1,29)}$=5.145, p<0.05).
Each dot represents the average value for one animal. n=7-9. Mean ± SEM.

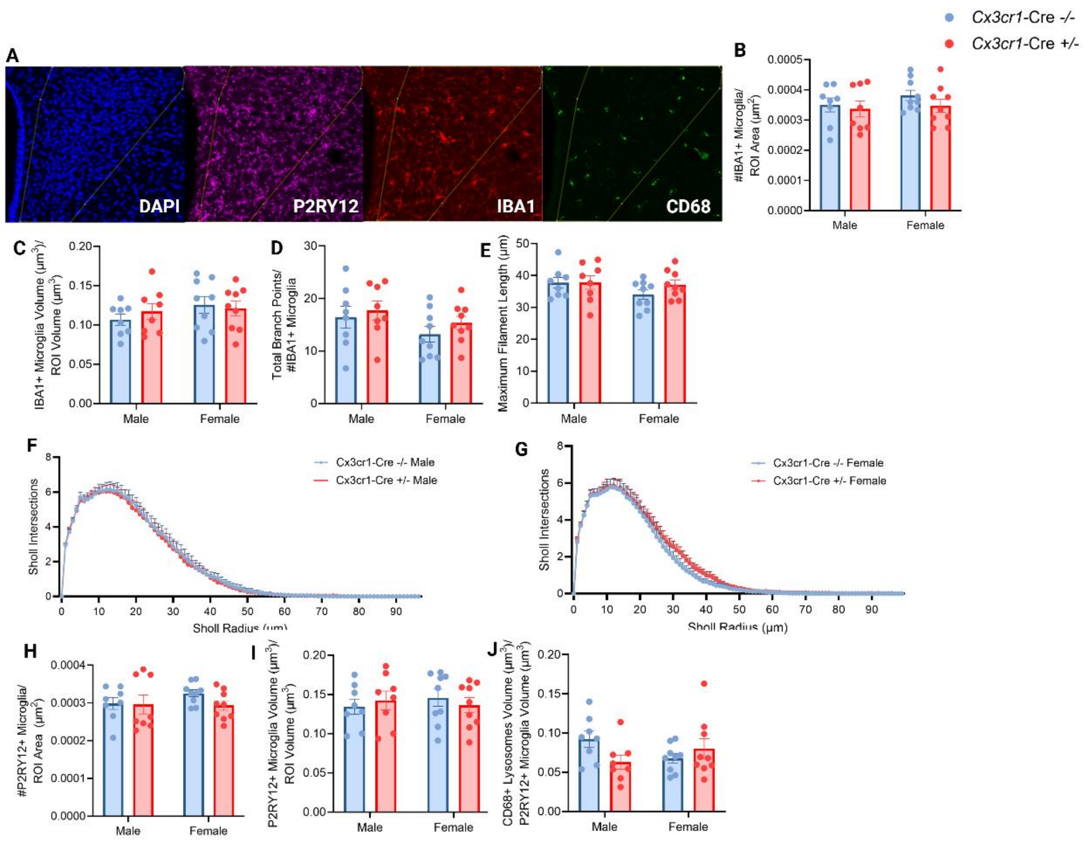

**Figure S12. Effects of constitutive CX3CR1-BAC-Cre expression on microglia in the adult PVN.**

A. Representative images showing DAPI, P2RY12, IBA1 and CD68 staining in the PVN.
B. Quantification of IBA1+ microglia density.
C. Quantification of IBA1+ microglia volume.
D. (D-G) IBA1 Sholl analysis metrics. Total branch points of IBA1+ microglia.
E. Maximum filament length of IBA1+ microglia.
F. Number of Sholl intersections for each radius of IBA1+ microglia of male mice.
G. Number of Sholl intersections for each radius of IBA1+ microglia of female mice.
H. Quantification of P2RY12+ microglia density.
I. Quantification of P2RY12+ microglia volume.
J. Quantification of CD68+ microglial lysosome volume relative to the P2RY12+ microglial volume.

Each dot represents the average value for one animal. n=7-9. *$p<0.05$. Mean ± SEM.

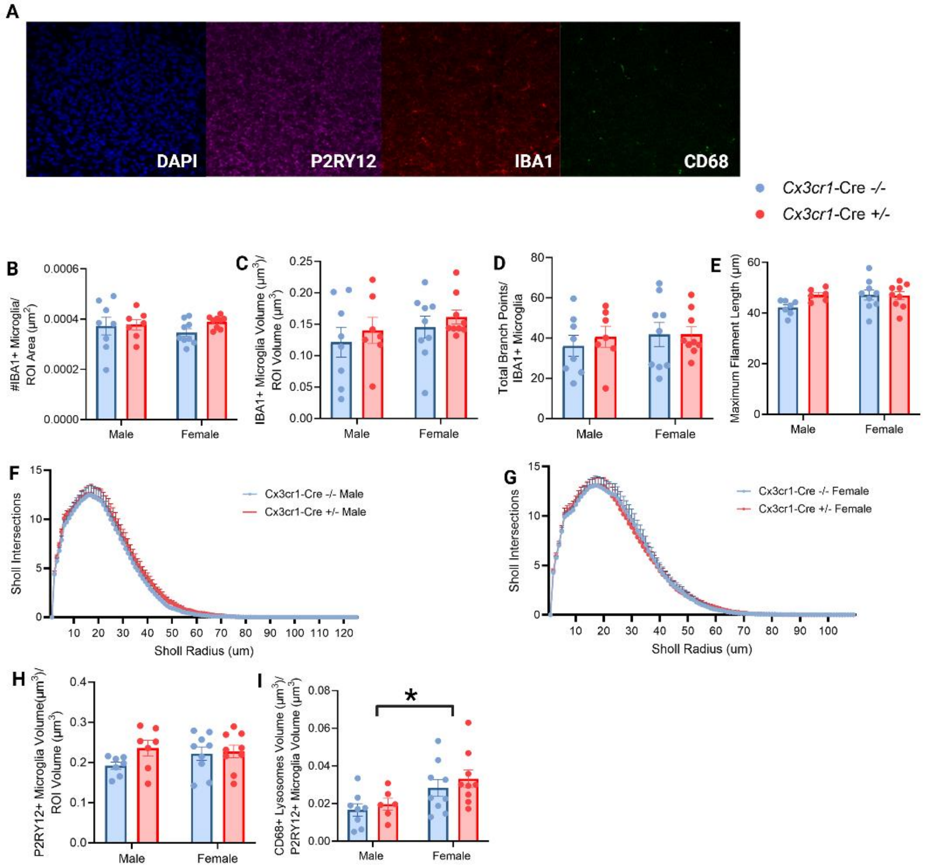


**Figure S13. Effects of constitutive CX3CR1-BAC-Cre expression on microglia in the adult Amygdala.**

A. Representative images showing DAPI, P2RY12, IBA1 and CD68 staining in the Amygdala.
B. Quantification of IBA1+ microglia density.
C. Quantification of IBA1+ microglia volume.
D. (D-G) Sholl analysis metrics. Total branch points of IBA1+ microglia.
E. Maximum filament length of IBA1+ microglia.
F. Number of Sholl intersections for each radius of IBA1+ microglia of male mice.
G. Number of Sholl intersections for each radius of IBA1+ microglia of female mice.
H. Quantification of P2RY12+ microglia volume.
I. Quantification of CD68+ microglial lysosome volume relative to the P2RY12+ microglial volume (main effect of Sex, $F_{(1,27)}$=5.167, p<0.05).

Each dot represents the average value for one animal. n=7-9. *p<0.05. Mean ± SEM.

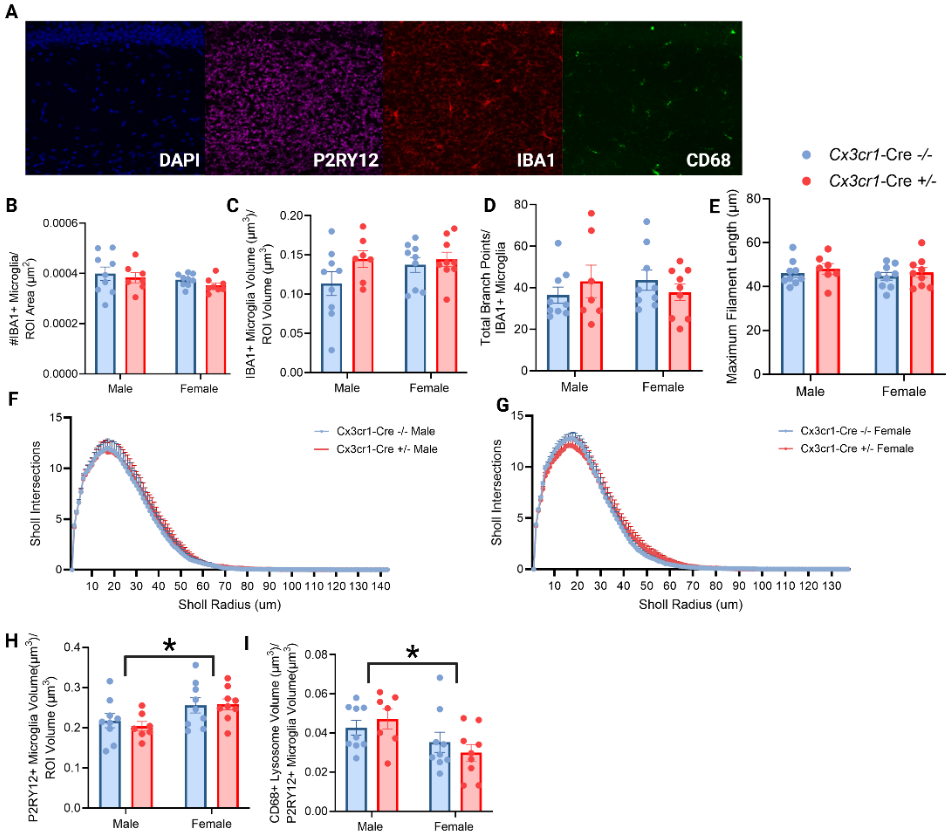


**Figure S14. Effects of constitutive CX3CR1-BAC-Cre expression on microglia in the adult CA1.**

A. Representative images showing DAPI, P2RY12, IBA1 and CD68 staining in the CA1.
B. Quantification of IBA1+ microglia density.
C. Quantification of IBA1+ microglia volume.
D. (D-G) Sholl analysis metrics. Total branch points of IBA1+ microglia.
E. Maximum filament length of IBA1+ microglia.
F. Number of Sholl intersections for each radius of IBA1+ microglia of male mice.
G. Number of Sholl intersections for each radius of IBA1+ microglia of female mice.
H. Quantification of P2RY12+ microglia volume (main effect of Sex, $F_{(1,30)}=7.977$, $p<0.05$).
I. Quantification of CD68+ microglial lysosome volume relative to the P2RY12+ microglial volume (main effect of Sex, $F_{(1,30)}=7.194$, $p<0.05$).
Each dot represents the average value for one animal. n=7-9. *$p<0.05$. Mean ± SEM.

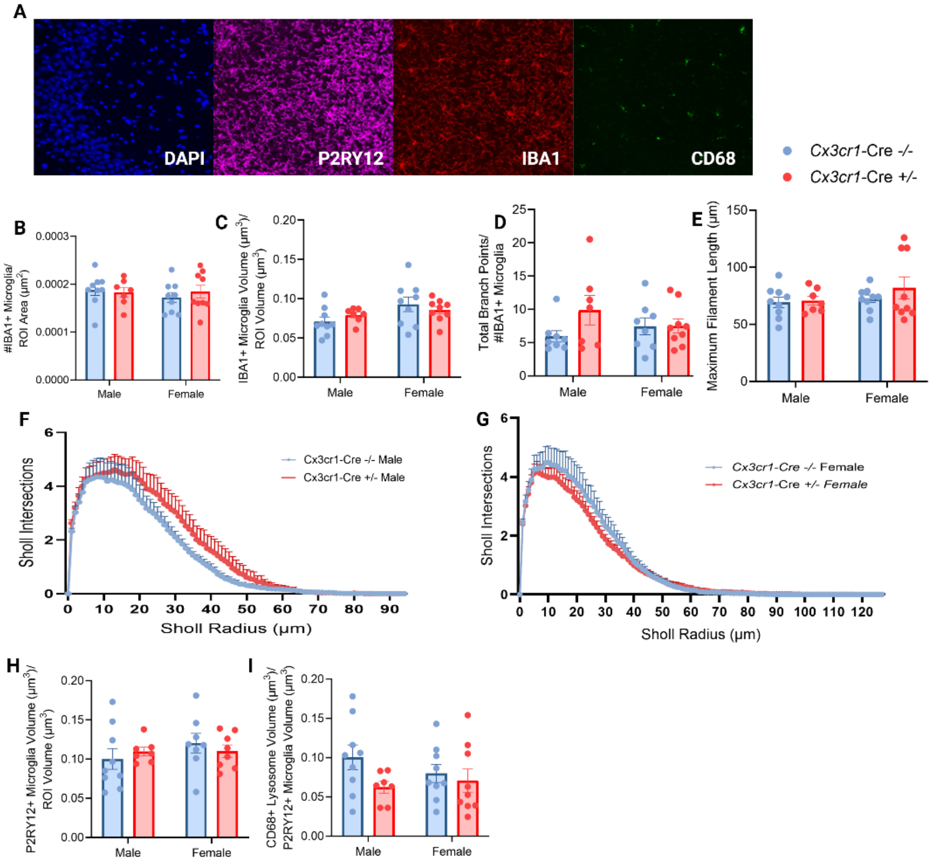


**Figure S15. Effects of constitutive CX3CR1-BAC-Cre expression on microglia in the adult CA3.**

A. Representative images showing DAPI, P2RY12, IBA1 and CD68 staining in the CA3.
B. Quantification of IBA1+ microglia density.
C. Quantification of IBA1+ microglia volume.
D. (D-G) Sholl analysis metrics. Total branch points of IBA1+ microglia.
E. Maximum filament length of IBA1+ microglia.
F. Number of Sholl intersections for each radius of IBA1+ microglia of male mice.
G. Number of Sholl intersections for each radius of IBA1+ microglia of female mice.
H. Quantification of P2RY12+ microglia volume.
I. Quantification of CD68+ microglial lysosome volume relative to the P2RY12+ microglial volume.

Each dot represents the average value for one animal. n=7-9. *$p<0.05$. Mean ± SEM.

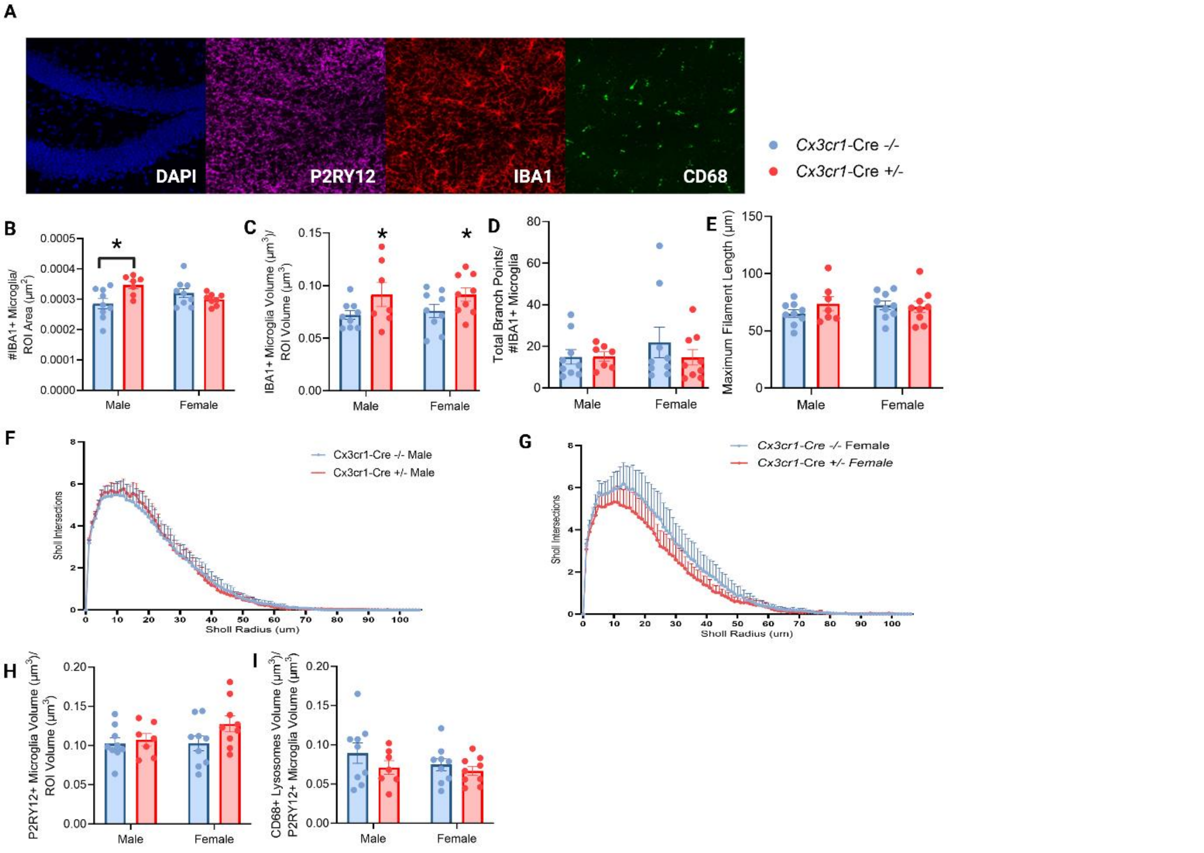


**Figure S16. Effects of constitutive CX3CR1-BAC-Cre expression on microglia in the adult DG.**

A. Representative images showing DAPI, P2RY12, IBA1 and CD68 staining in the DG.
B. Quantification of IBA1+ microglia density (interaction Sex x Genotype, $F_{(1, 29)}$=8.964, p=<0.05; Šídák, p<0.05).
C. Quantification of IBA1+ microglia volume (main effect of Genotype, $F_{(1,30)}$=6.128, p<0.05).
D. (C-F) Sholl analysis metrics. Total branch points of IBA1+ microglia.
E. Maximum filament length of IBA1+ microglia.
F. Number of Sholl intersections for each radius of IBA1+ microglia of male mice.
G. Number of Sholl intersections for each radius of IBA1+ microglia of female mice.
H. Quantification of P2RY12+ microglia volume.
I. Quantification of CD68+ microglial lysosome volume relative to the P2RY12+ microglial volume.

Each dot represents the average value for one animal. n=7-9. *p<0.05. Mean ± SEM.
